# SARS-CoV-2 gene content and COVID-19 mutation impact by comparing 44 Sarbecovirus genomes

**DOI:** 10.1101/2020.06.02.130955

**Authors:** Irwin Jungreis, Rachel Sealfon, Manolis Kellis

## Abstract

Despite its overwhelming clinical importance, the SARS-CoV-2 gene set remains unresolved, hindering dissection of COVID-19 biology. Here, we use comparative genomics to provide a high-confidence protein-coding gene set, characterize protein-level and nucleotide-level evolutionary constraint, and prioritize functional mutations from the ongoing COVID-19 pandemic. We select 44 complete Sarbecovirus genomes at evolutionary distances ideally-suited for protein-coding and non-coding element identification, create whole-genome alignments, and quantify protein-coding evolutionary signatures and overlapping constraint. We find strong protein-coding signatures for all named genes and for 3a, 6, 7a, 7b, 8, 9b, and also ORF3c, a novel alternate-frame gene. By contrast, ORF10, and overlapping-ORFs 9c, 3b, and 3d lack protein-coding signatures or convincing experimental evidence and are not protein-coding. Furthermore, we show no other protein-coding genes remain to be discovered. Cross-strain and within-strain evolutionary pressures largely agree at the gene, amino-acid, and nucleotide levels, with some notable exceptions, including fewer-than-expected mutations in nsp3 and Spike subunit S1, and more-than-expected mutations in Nucleocapsid. The latter also shows a cluster of amino-acid-changing variants in otherwise-conserved residues in a predicted B-cell epitope, which may indicate positive selection for immune avoidance. Several Spike-protein mutations, including D614G, which has been associated with increased transmission, disrupt otherwise-perfectly-conserved amino acids, and could be novel adaptations to human hosts. The resulting high-confidence gene set and evolutionary-history annotations provide valuable resources and insights on COVID-19 biology, mutations, and evolution.

## Introduction

SARS-CoV-2, the virus responsible for COVID-19^1^, is a betacoronavirus in the subgenus Sarbecovirus, which also includes SARS-CoV, responsible for the 2003 severe acute respiratory syndrome (SARS) outbreak. Its large 29,903-nucleotide positive-strand RNA genome encodes ∼30 known and hypothetical mature proteins (**Fig. 1a, Fig. 2, Extended Data Fig. 1**). Despite SARS-CoV-2’s extreme medical importance, its gene content remains surprisingly unresolved, with several hypothetical open reading frames (ORFs) whose function or even protein-coding status is unknown. Moreover, no systematic resource exists for interpreting the functional impact of SARS-CoV-2 mutations and prioritizing candidate drivers that may underlie phenotypic differences between strains.

**Figure 1.**
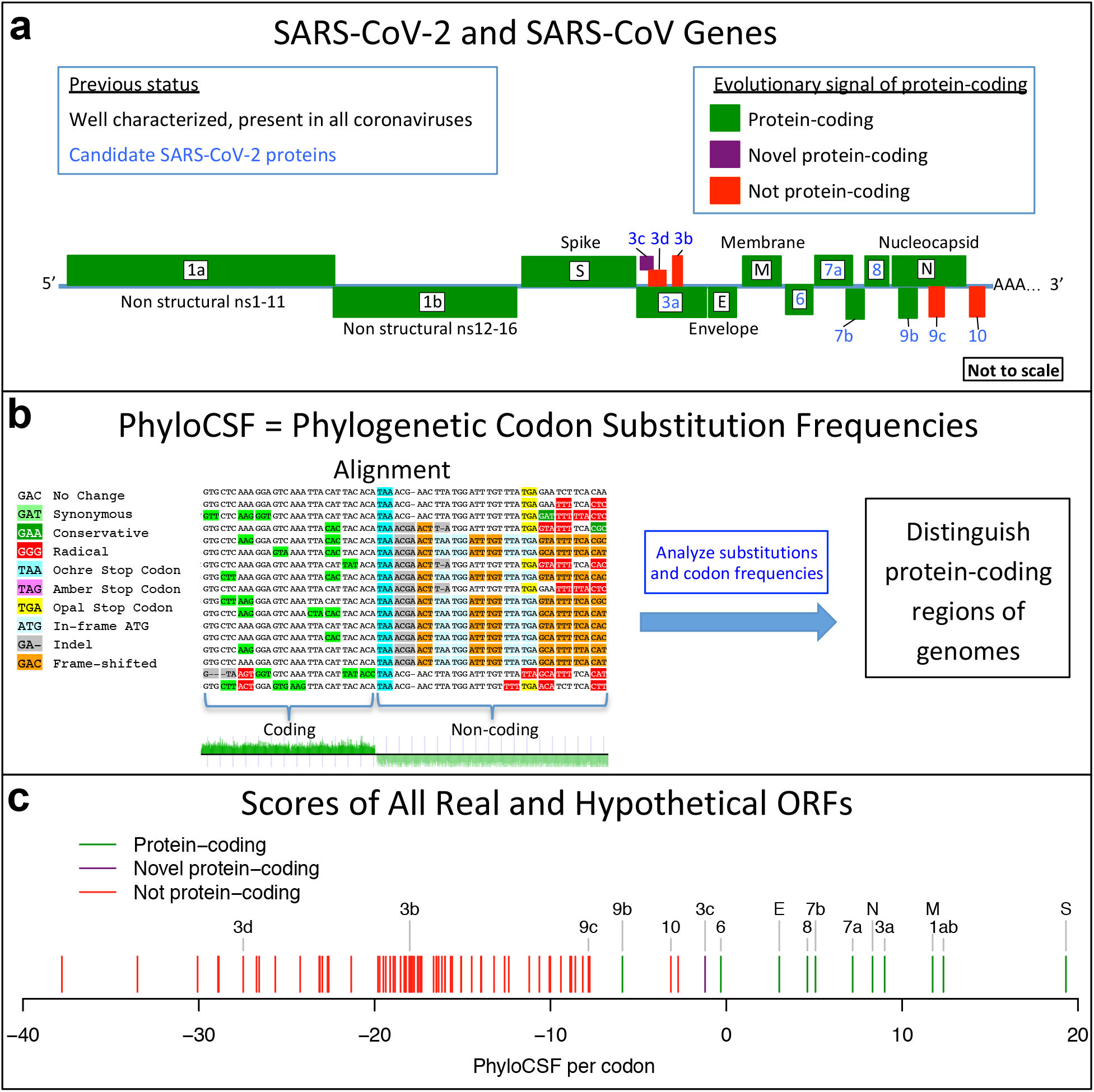
Overview. **a.** Previously annotated named (black font) and unnamed or proposed (blue font) SARS-CoV-2 genes, with confirmed protein-coding (green), rejected (red), or novel protein-coding (purple) classification, using evolutionary and experimental evidence. **b.** Phylogenetic Codon Substitution Frequencies (PhyloCSF) scores distinguish protein-coding (left) vs. non-coding (right) using evolutionary signatures, including distinct frequencies of amino-acid-preserving (green) vs. amino-acid-disruptive (red) substitutions, and stop codons (cyan/magenta/yellow) in frame-specific alignments, and additional features. **c.** PhyloCSF score (x-axis) for all confirmed (green) and rejected (red) ORFs, showing annotated/hypothetical/novel (labeled) and all AUG-initiated ≥25-codons-long locally-maximal ORFs (unlabelled). Novel ORF3c (purple) clusters with protein-coding. Only-modestly-negative ORF9c/ORF10 scores are artifacts of score compression in high-nucleotide-constraint regions, and substantially drop when nucleotide-conservation-scaled (see **Extended Data Fig. 8**).

**Figure 2.**
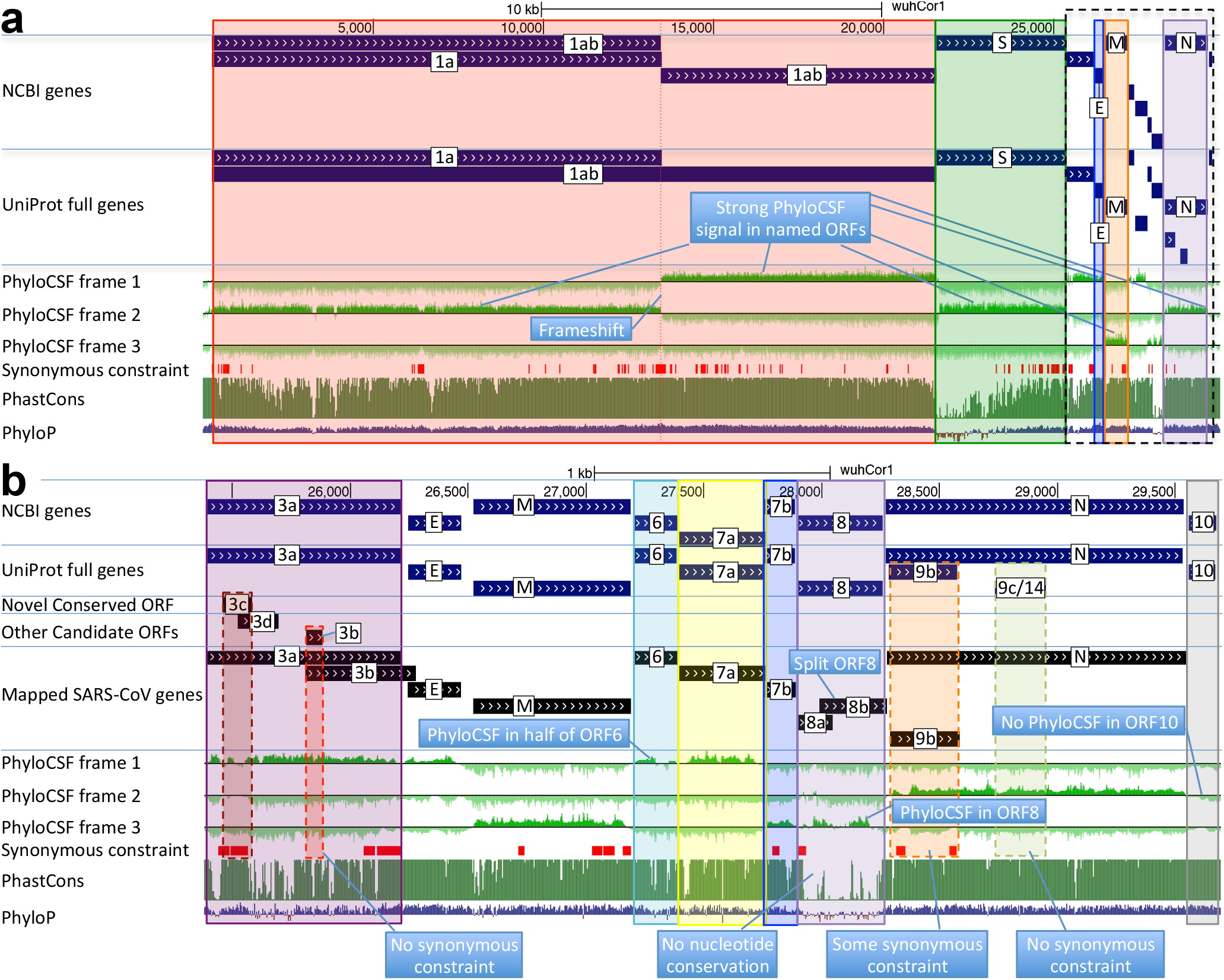
Genome-wide protein-coding signatures. SARS-CoV-2 NCBI/UniProt genes (blue), unannotated proposed genes and mapped SARS-CoV genes (black, panel b only), frame-specific protein-coding PhyloCSF scores (green), Synonymous Constraint Elements (SCEs) (red), and phastCons/phyloP nucleotide-level constraint (green/blue/red) across genomic coordinates (x-axis) for entire genome (panel a) and final 4-kb subset (panel b, dashed black box), highlighting (light blue boxes): **(a)** strong protein-coding signal in correct frame for each named gene; conservation-signal frame-change at programmed frameshift site; strong protein-coding signal throughout S despite lack of nucleotide conservation in S1; **(b)** unambiguous and frame-specific protein-coding signal for unnamed ORFs 3a (despite only partial nucleotide conservation), 7a, 7b, and 8 (despite lack of nucleotide conservation); clear protein-coding signal in first half and last quarter of ORF6; no protein-coding signal for 10 (despite high nucleotide conservation); synonymous constraint (red) in novel-ORF 3c and confirmed-ORF 9b; no synonymous constraint in rejected ORFs 9c, 3b, 3d.

A large open reading frame spans two thirds of the genome, and results in non-structural proteins nsp1-nsp10 and nsp12-nsp16 when an internal programmed translational frameshift^2^ occurs (ORF1ab), or nsp1-11 otherwise (ORF1a) with translation terminating four codons past the frameshift site. ORF1ab encodes Pol (polymerase, RNA-dependent replication), Hel (helicase), ExoN (exonuclease, proofreading), 3CL-PRO (polyprotein cleavage), and other proteins involved in host-cell suppression, immune suppression, and diverse viral functions (Supplementary Table S2).

The last third of the genome encodes named proteins S (Spike surface glycoprotein), composed of S1 (viral attachment to host-cell ACE2 receptor) and S2 (membrane fusion, viral entry), E (Envelope protein), M (Membrane glycoprotein), and N (Nucleocapsid, RNA genome packaging), which are present in all coronaviruses, and several unnamed proteins. Their host-cell translation requires subgenomic RNAs of varying lengths, such that each functional ORF is first (or early) on its own transcript^3^. This results from positive-to-negative transcription from the 3’ end to a transcription-regulatory sequence (TRS), looping to a common 5’ leader, followed by negative-to-positive transcription^4^.

The remaining unnamed ORFs are Sarbecovirus-specific and subject to disagreement on which encode functional proteins (Supplementary Table S2). NCBI annotates SARS-CoV-2 (NC_045512.2) with 3a, 6, 7a, 7b, 8, and 10. UniProt also annotates 9b and 9c (which they name 14), both overlapping N (in an alternate frame). The paper introducing SARS-CoV-2 also shows 3b (which overlaps 3a in SARS-CoV but is truncated in SARS-CoV-2, with several in-frame stop codons)^1^. Other publications^5–13^ include different subsets, use different names, or propose additional ORFs (including 3c and 3d overlapping 3a). NCBI annotates SARS-CoV (NC_004718.3) orthologs of 3a, 6, 7a, 7b, and 9b, but 8 is split into 8a and 8b, 3b is included, and neither 9c nor 10 are included. (ORF nomenclature details in **Supplementary Text S1**.)

High-throughput experiments provide some evidence on SARS-CoV-2 gene content, though they sometimes disagree, cannot prove non-functionality of non-detected ORFs (as they only capture specific conditions), and cannot distinguish incidental transcriptional/translational events from selected function. Proteomics identified peptides for 1ab, S, 3a, M, 6, 7a, 8, N, and 9b, but not E, 3b, 7b, 9c, or 10^14,15^. Direct-RNA sequencing found subgenomic RNAs for a different subset: S, 3a, E, M, 6, 7a, 7b, 8, and N, but limited or no support for 3b, 3d, 3c, 9b, 9c, and 10^15–18^, with 3c, 7b^19^, and 9b possibly translated by leaky ribosome scanning from 3a, 7a, and N subgenomic RNAs, respectively. Ribosome profiling predicted translation of 1ab, S, 3a, E, M, 6, 7a, 7b, 8, N, and 10, and eleven alternate-frame ORFs (including 3c, 9b), but not ORFa 3d, 3b, or 9c^10^.

Here, we use comparative genomics of 44 Sarbecovirus strains to resolve the SARS-CoV-2 protein-coding gene set (**Fig. 1**), and to distinguish genetic variants more likely to have functional importance. We select 44 closely-related and complete coronavirus genomes, generate whole-genome alignments, evaluate protein-coding and nucleotide-level constraint, and annotate synonymously-constrained codons. We show that five hypothetical ORFs are not functional proteins and confirm protein-coding status for seven accessory ORFs, including novel alternate-frame ORF3c within 3a. We use protein-level and nucleotide-level inter-strain constraint to analyze 1875 mutations from 2544 pandemic isolates, show gene-level and codon-level agreement between within-strain and across-strain selective pressures, reveal recent adaptive acceleration for N and surprising deceleration for S1 and nsp3, and flag mutations disrupting evolutionarily-conserved positions that may represent novel adaptations to human hosts, including Spike D614G.

## Results

### Strain selection, alignment, constraint

We selected and aligned 44 complete Sarbecovirus genomes (SARS-CoV-2, SARS-CoV, and 42 bat-infecting strains, **Extended Data Fig. 2, Supplementary Table S1**) at evolutionary distances well-suited for identifying protein-coding genes and non-coding purifying selection, spanning ∼3 substitutions per 4-fold degenerate site on average (comparable to 29-mammals/12-flies projects^20,21^), and ranging from 1.2 (E) to 4.8 (nsp16) and higher (**Supplementary Table S2**). Betacoronaviruses outside Sarbecovirus (including MERS-CoV) are too distant (eg. no detectable homology across ORFs 6-7a-7b-8), and SARS-CoV-2/SARS-CoV isolates are too proximal for reliable evolutionary signatures.

To distinguish regions evolving under protein-coding constraint, we used their codon substitution patterns across Sarbecoviruses, quantified using codon-resolution PhyloCSF^22^ scores in all three reading frames, and smoothed using a hidden Markov model to create genome browser tracks^1,23,24^ (**Fig. 1b, Fig. 2**). We also computed gene-resolution PhyloCSF scores for each known protein and hypothetical ORF, and generated CodAlignView^25^ visualizations highlighting protein-coding vs. non-coding features for manual exploration of their alignments in all reading frames (**Fig. 1c, Supplementary Table S2**). These tools are widely-accepted standards for protein-coding gene annotation and for distinguishing protein-coding vs. non-coding genes in human and other species^20– 22,26–28^.

Beyond protein-coding constraint for amino-acid translation, we also evaluated nucleotide-level overlapping constraint within protein-coding regions indicative of dual-coding regions, RNA structures, RNA-binding protein sites, etc, using reduced synonymous-substitution rate estimated using FRESCo, which we previously developed and applied to viruses^29^ and human^30^. We annotated 1394 synonymously-constrained codons (14% of 9744, FDR=0.125) and defined 92 synonymous-constraint elements (SCEs) (covering 1555 codons), using 9-codon-resolution significantly-decreased synonymous rate relative to gene average^29,31^.

### Coding constraint on non-overlapping genes

As expected, E, M, N, S2, nsp1-nsp10, and nsp12-nsp16 showed clear protein-coding constraint (**Supplementary Table S2**), with a change in constrained reading frame at the known programmed frameshift (**Fig. 2, Extended Data Fig. 1**). Beyond its first 9 codons that match Pol, the 13-codon nsp11 showed no nucleotide changes in Sarbecovirus, but stop-codon gain/loss across betacoronaviruses indicates it is not separately functional (**Supplementary Fig. S1**).

S1 shows extremely rapid nucleotide evolution (near-zero phyloP^32^ and phastCons^33^) but strong PhyloCSF scores, indicating unambiguous protein-coding evolution and highlighting the power of PhyloCSF to recognize protein-coding evolutionary signatures despite rapid nucleotide evolution.

ORFs 3a, 7a, 7b, and 8 show clear positive PhyloCSF scores, indicating conserved protein-coding regions functional at the amino acid level (**Fig. 2b**). The first half and last quarter of ORF6 show strong PhyloCSF signal, indicating that it encodes a functional protein, despite a less-constrained intermediate portion, and an overall near-zero average score per codon (−0.3, **Fig. 1c**).

ORF8 shows near-zero nucleotide-level conservation (phyloP/phasCons), lacks well-established functions, and was split into 8a/8b in SARS-CoV, suggesting at first glance that it might be non-functional. However, it shows strongly-positive protein-coding PhyloCSF score (4.61/codon), and long stretches of strong protein-coding constraint, indicating unambiguous protein-coding function. Its high nucleotide-level rate is inflated by past recombination, but remains high even using an ORF8-specific phylogeny (**Supplementary Fig. S2**).

By contrast, ORF10 shows no protein-coding constraint anywhere along its length, contains in-frame stop codons in all but four Sarbecoviruses truncating the last third of its already-short length (38 amino acids), includes a frame-shifting deletion in one of those four strains, and shows near-perfect nucleotide-level conservation (phyloP/phastCons) extending beyond the ORF on both sides, indicating it is not protein-coding but instead has non-coding functions (**Fig. 2b, Extended Data Fig. 3a**). (This region overlaps the 3’-UTR pseudoknot RNA structure^34^ involved in RNA synthesis, providing a likely explanation for its high nucleotide-level constraint). Moreover, ribosome footprints in the region occur in an overlapping upstream ORF or in a truncated ORF rather than uniquely in ORF10, consistent with incidental-initiation events rather than functional translation (**Extended Data Fig. 3b**), and previously-used comparative evidence for protein-coding function ignored a frameshifting deletion and was insufficiently-powered (**Extended Data Fig. 3c**).

### N-overlapping ORF 9b is coding, 9c is not

Evolutionary evidence for/against overlapping ORFs is harder to resolve, as protein-coding signatures in the primary reading frame heavily influence scores in alternate frames: they skew the signal as protein-preserving mutations in one frame are typically protein-disruptive in the other, and they compress the signal as there are fewer substitutions. However, their dual-coding nature leads to a depletion of synonymous substitutions in the primary ORF localized over the overlapping segment, resulting in a strong signal of overlapping-constraint^29–31^, used next to investigate ORFs 9c and 9b overlapping N.

The 73-amino-acid-long ORF9c/ORF14 shows no localized synonymous constraint in N (**Fig. 3**), calling its protein-coding status into question. Moreover, its start codon is lost in one strain, most strains have a three-codons-earlier stop (**Extended Data Fig. 4**), its start codon is 460 nucleotides after N’s with 9 intervening AUG codons (thus unlikely to be translated via leaky ribosome scanning), direct-RNA sequencing found no ORF9c-specific subgenomic RNAs^16–18^ (and no TRS is appropriately positioned to create one), shows no ribosome footprint^10^ or proteomics^14,15^ evidence, and many SARS-CoV-2 isolates^35^ contain stop-introducing mutations^7^. We conclude ORF9c does not encode a functional protein.

**Figure 3.**
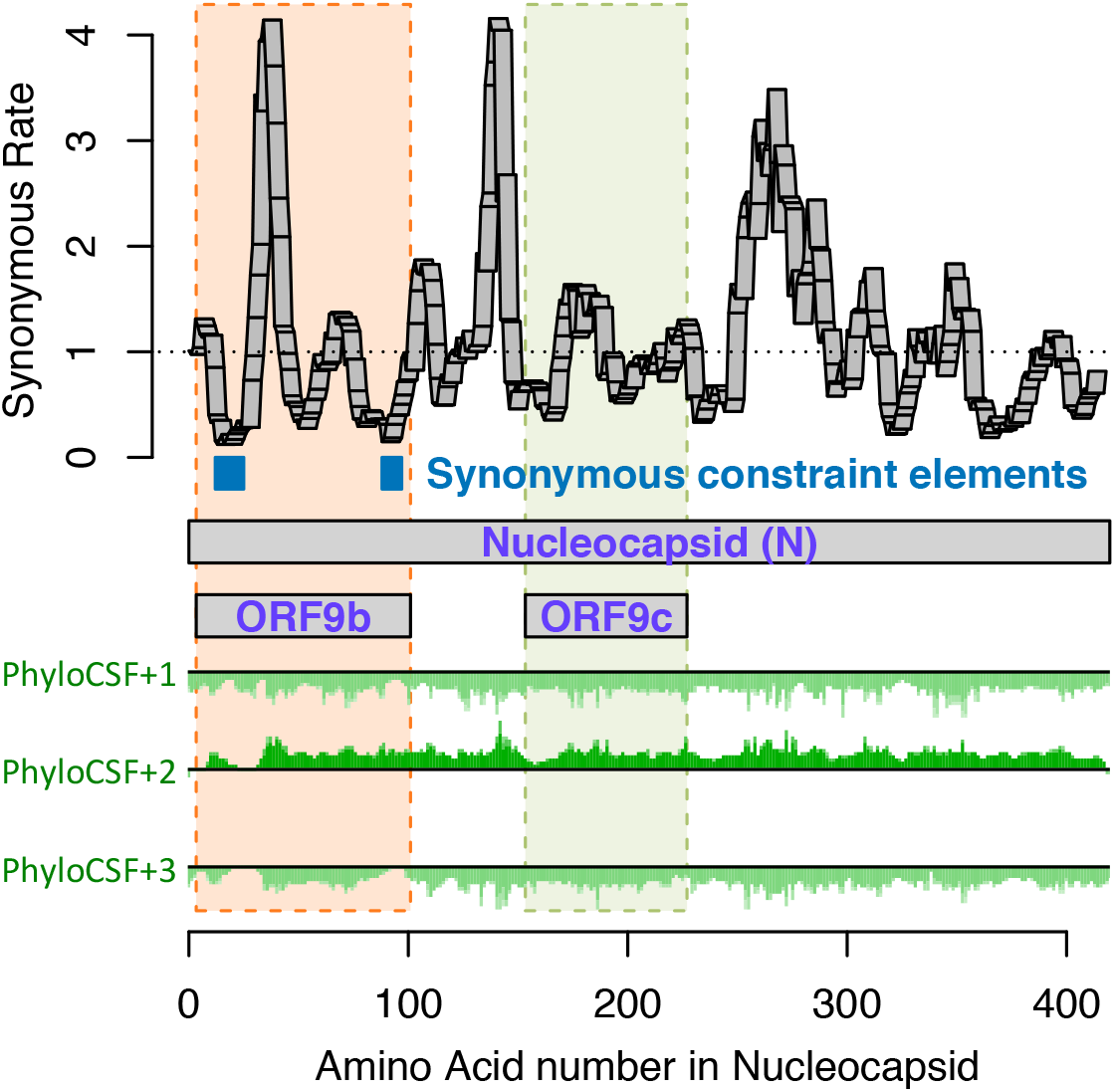
Synonymous constraint in Nucleocapsid overlaps 9b but not 9c/14. Synonymous substitution rate in 9-codon windows (y-axis) across N (x-axis), normalized to gene-wide average (dotted black line). Synonymous constraint elements (blue) expected for dual-coding constraint localize in overlapping ORF9b (dashed orange rectangle) indicating it is protein-coding, but not 9c (dashed purple rectangle) indicating it is not protein-coding. PhyloCSF protein-coding signal (green) in frame3 (encoding 9b and 9c/14) remains strongly negative throughout the length of 9c/14 (green box), indicating 9c/14 is non-coding, but rises to near-zero values for two regions of 9b, indicating protein-coding selection, while PhyloCSF signal frame 2 (encoding N) remains consistently high throughout the length of ORF9c.

The 97-amino-acid-long ORF9b shows high amino-acid substitution rate in its central portion but significant localized synonymous constraint in N for its start and end regions (**Fig. 3**), even relative to the overall low synonymous rate of N, consistent with dual-coding functions. Moreover, its start and stop codons are perfectly conserved and its 97 codons are stop-free in all Sarbecoviruses. Its Kozak context is stronger than N’s and perfectly-conserved and its start codon is only 10 nucleotides downstream of N’s, allowing it to be translated from N’s subgenomic RNA via leaky scanning (**Extended Data Fig. 5**). ORF9b’s negative PhyloCSF score is consistent with dual-coding signal biases. ORF9b also has proteomics support^15,36,37^ (including evidence of viral-RNA binding^38^), and alternate-frame translation support by ribosome profiling^10^. In SARS-CoV, ORF9b protein (and antibodies to it) was detected in SARS patients^39,40^, localized in mitochondria, and interfered with host cell antiviral response when overexpressed^41^. We conclude ORF9b encodes a conserved functional protein with rapidly-changing portions.

### ORF3c is a novel functional protein

We next searched for additional protein-coding genes by computing PhyloCSF scores for all 67 hypothetical non-NCBI-annotated AUG-to-stop SARS-CoV-2 ORFs ≥25 codons long that are not contained in a longer same-frame ORF (locally-maximal). None had positive PhyloCSF scores, but some may be coding as overlapping-ORF scores are reduced by alternative-frame protein-coding constraint, so we investigated near-zero top candidates for evidence of localized synonymous constraint, start/stop-codon conservation, and absence of in-frame stops or frameshifting indels.

The highest-scoring candidate, which we call ORF3c, overlaps ORF3a near its start (**Fig. 4**), with 38 of its 41 codons overlapping synonymous constraint elements in ORF3a, localized nearly-perfectly on the dual-coding region. Despite the score biases of dual-coding regions, ORF3c has PhyloCSF score closer to non-overlapping protein-coding ORFs than to hypothetical non-coding ORFs (**Fig. 1c**), indicating Sarbecovirus selection for protein-coding function. Strikingly, ORF3c also has many synonymous substitutions that are non-synonymous in ORF3a, indicating ORF3c may be an equally-strong driver of constraint in the dual-coding region (both frames show similar scores in the dual-coding region). ORF3c also has conserved start and stop codons except for near-cognate GUG start in one strain and a one-codon extension in SARS-CoV-2 and RaTG13, with no in-frame stop codons or indels. We conclude ORF3c encodes a functional, conserved protein.

**Figure 4.**
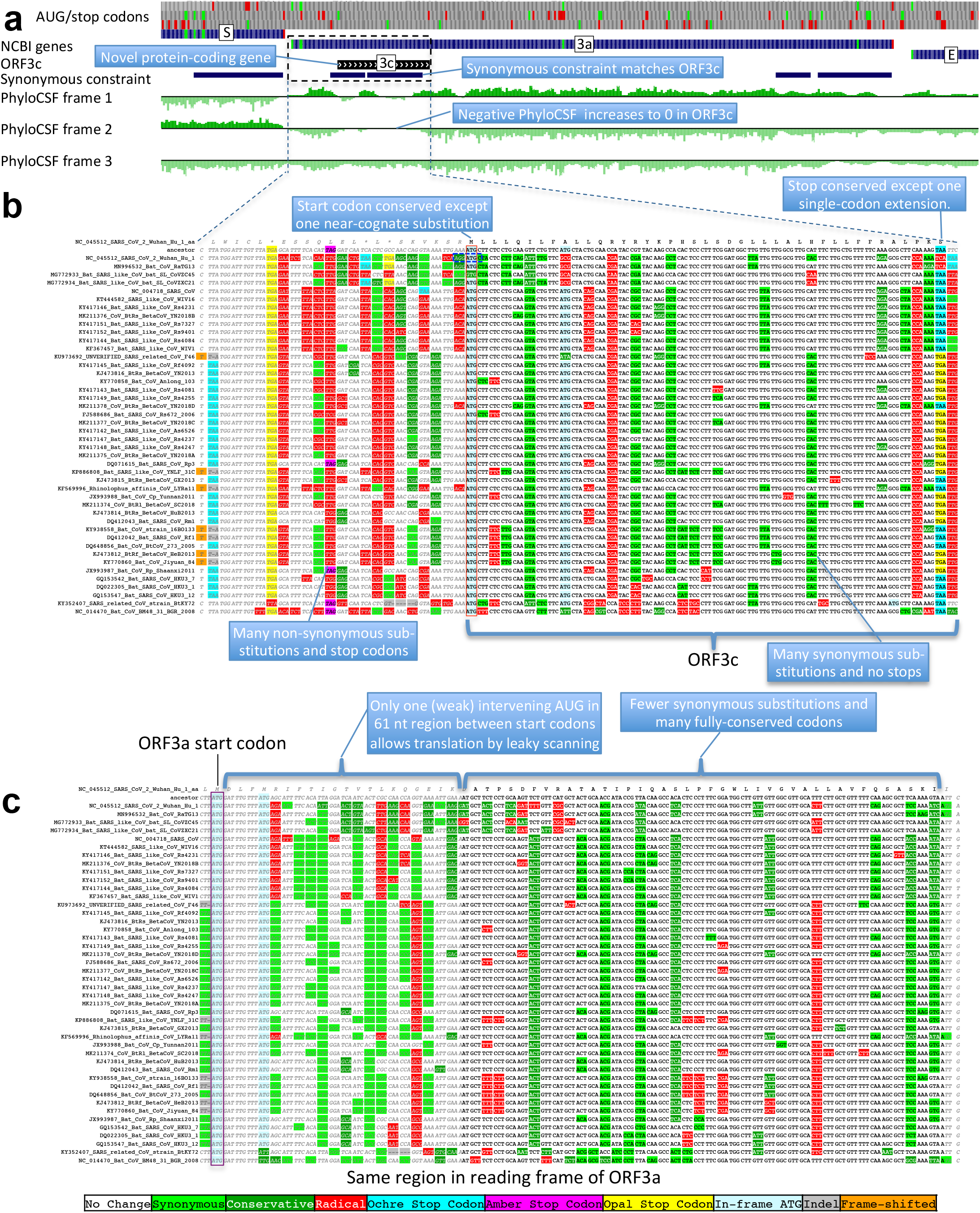
Novel gene 3c overlapping 3a is protein-coding. **a.** Synonymous-constraint elements (blue) match nearly-perfectly 41-codon ORFc dual-coding region boundaries (black), and protein-coding evolutionary signatures (green) switch between frame 1 and 2 (rows) in the dual-coding region, with frame-2 signal (negative flanking ORF3c) increasing to near-zero, and frame-1 signal (high flanking the dual-coding region) dropping to near-zero. **b**,**c**. Codon-resolution evolutionary signatures (colors, CodAlignView^25^) annotating genomic alignment (letters) spanning ORF3a start and dual-coding region, in frame-1 (top) and frame-2 (bottom), highlighting (blue boxes): (**b**, frame-2, ORF3c) radical codon substitutions (red) and stop codons (yellow, magenta, cyan) prior to ORF3c start; synonymous (light green) and conservative (dark green) substitutions in ORF3c; ORF3c’s start codon is conserved, except in one strain (row 4) with near-cognate GUG; ORF3c’s stop codon is conserved except for one-codon extension in two strains (rows 2-3); no intermediate stop codons in ORF3c; (**c**, frame-1, ORF3a) abundant synonymous and conservative substitutions in 3a prior to dual-coding region; increase in fully-conserved codons (white) over dual-coding region. Short interval (61nt) with only one weak-Kozak-context intervening start codon indicates ORF3c may be translated from ORF3a’s subgenomic RNA via leaky scanning.

Previous studies proposed four ORFs overlapping 3a^6,8–13^: 3c (41 codons), 3d (57 codons), 3b (22 codons, a truncated ortholog of SARS-CoV ORF3b), and a subset of 3d (33 codons). ORF3c was proposed using synonymous constraint across 6 closely-related strains^8^ and a broader set of Sarbecoviruses^9^, although on its own such evidence could also stem from other overlapping functional elements (and is abundant in SARS-CoV-2 even outside dual-coding regions), and using ribosome footprinting^10^, although such signal can also result from incidental, non-functional translation (and the other 8 such candidates lacked any conservation); it was predicted to contain a viroporin-like transmembrane domain^8^ and to be translated via leaky scanning^9^. The other three ORF3a-overlapping candidates are not conserved and show variable length, premature stop codons, and other evidence indicating they are not protein-coding (**Extended Data Fig. 6, Extended Data Fig. 7, Supplementary Text S2**).

We examined all next-best-scoring candidates, and expanded the search to include shorter ORFs, near-cognate start codons, non-locally-maximal ORFs, and ORFs on the negative strand, but found no other convincing candidates (**Supplementary Text S3, Supplementary Fig. S4**), concluding our protein-coding gene catalog is complete.

### A new reference gene set for SARS-CoV-2

Altogether, our revised reference gene set consists of 1a, 1ab, S, 3a, 3c, E, M, 6, 7a, 7b, 8, N, and 9b, including novel ORF 3c and previously-ambiguous 9b, and excluding 3b, 3d, 9c, and 10. These genes are unambiguously translated into conserved functional proteins across Sarbecoviruses, and our decisions are supported by a wealth of experimental evidence^10,14–18^, including subgenomic RNAs^15–18^ (or leaky scanning), ribosome profiling^10^, and proteomics experiments^14,15^(Supplementary Text S4). This high-confidence reference gene set can form the basis for understanding viral biology and the functional roles of pandemic mutations (**Supplementary Text S5**).

### Sarbecovirus conservation informs SARS-CoV-2 variant impact

We next used the evolutionary history of each codon across Sarbecoviruses to annotate 1875 single-nucleotide variants (SNVs) across 2544 SARS-CoV-2 isolates sequenced during the current COVID-19 pandemic, including 1142 amino-acid-changing (missense), 628 amino-acid-preserving (synonymous), and 104 non-coding substitutions (**Supplementary Table S3**).

We classified all amino acid positions as “conserved” (no change in any of the 44 Sarbecovirus genomes) or “non-conserved/changed” (at least one change) for each of the mature proteins and hypothetical ORFs (**Supplementary Table S2**), a definition independent of the phylogenetic tree, and thus resilient to recombination events common in coronavirus phylogenies^42^.

### Within-strain vs cross-strains evolution

The fraction of changed amino acids varied greatly across ORFs (17%-80%, **Fig. 5a, x-axis**), indicating dramatically different evolutionary pressures. Unnamed accessory ORFs had more changed amino acids (average 57%) than named and well-characterized ORFs (average 28%). ORF1ab mature proteins varied from 57% changed (nsp2) to <17% (3CL-PRO, Pol, Hel, ExoN, nsp7-10) and Spike subunits from 61% changed (S1) to 25% (S2).

**Figure 5.**
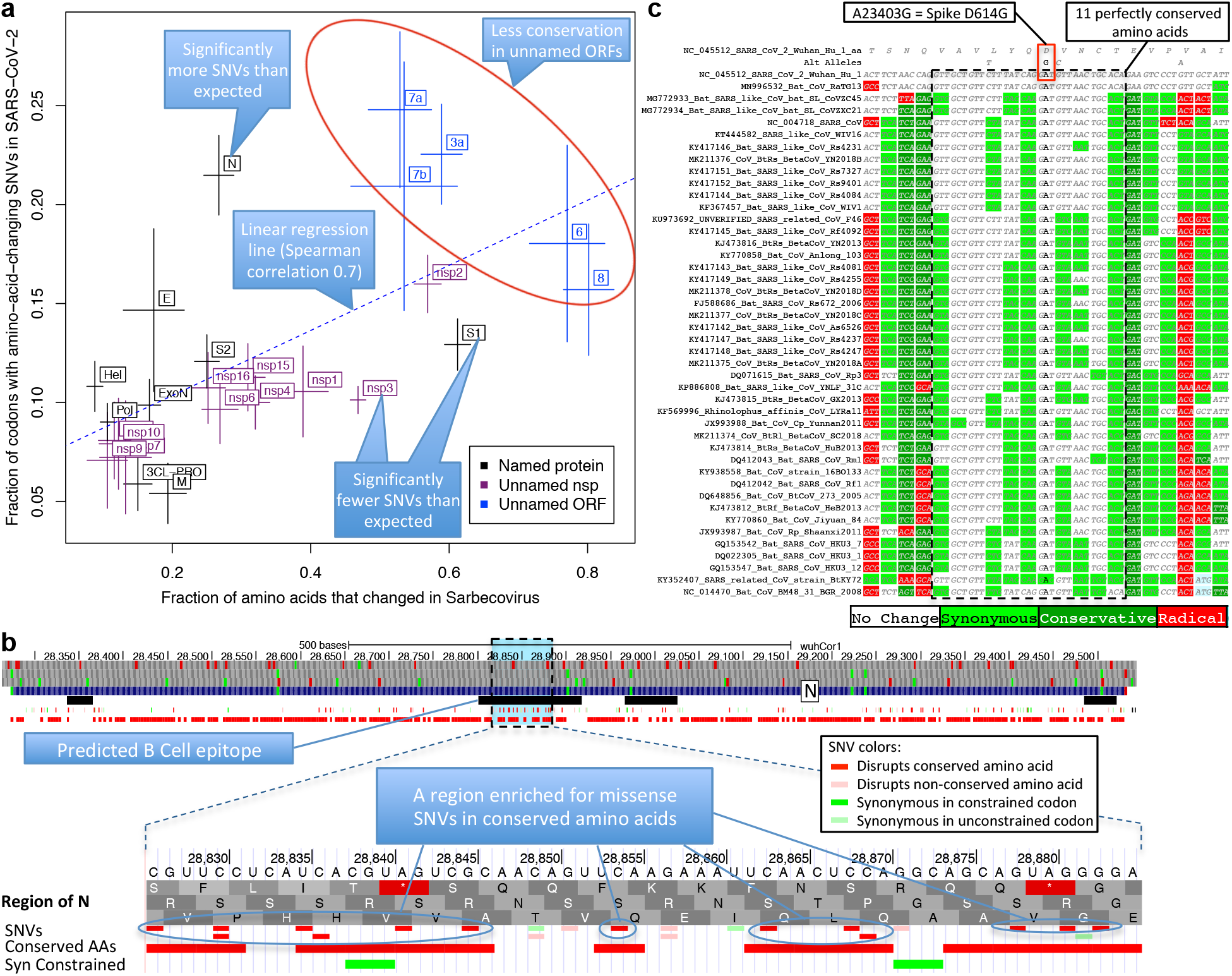
Within-strain variation vs. inter-strain divergence. **a. Gene-level comparison.** Long-term inter-strain evolutionary divergence (x-axis) and short-term within-strain variation (y-axis) show strong agreement (linear regression dotted line, Spearman-correlation=0.70) across mature proteins (crosses, denoting standard error of mean on each axis), indicating that Sarbecovirus-clade selective pressures persist in the current pandemic. Well-characterized genes (black) show fewer changes in both timescales (bottom left) and less-well-characterized ORFs (blue) show more in both (top right). Significantly-deviating exceptions are: nsp3 and S1 (bottom right) showing significantly-fewer amino-acid-changing SNVs than expected from their cross-Sarbecovirus rapid evolution, and N (top left), showing significantly-more, possibly due to accelerated evolution in the current pandemic. **b. Rapidly-evolving Nucleocapsid region.** Top: Nucleocapsid context showing B-cell epitope predictions (black, “IEDB Predictions” track), and our annotation track-hub showing: conserved amino acids (red blocks), synonymously-constrained codons (green blocks), and SNV classification (colored tick-marks) as conserved/non-conserved (dark/light) and missense/synonymous (red/green); top 3 tracks show AUG codons (green) and stop codons (red) in three frames. Bottom: Focus on 20-amino-acid region R185-G204 (dotted box) in predicted B-cell epitope (black) significantly-enriched for amino-acid-changing variants (red) disrupting perfectly-conserved residues, indicative of positive selection in SARS-CoV-2 for immune system avoidance. **c. Spike D614G evolutionary context.** Sarbecovirus alignment (text) surrounding Spike D614G amino-acid-changing SNV, which rose in frequency in multiple geographic locations suggesting increased transmissibility. This A-to-G SNV disrupts a perfectly-conserved nucleotide (bold font, A-to-G), which disrupts a perfectly-conserved amino-acid (red box, D-to-G), in a perfectly-conserved 11-amino-acid region (dotted black box, light-green=synonymous-substitutions) across bat-host Sarbecoviruses, indicating D614G represents a human-host-adaptive mutation.

Faster-evolving proteins across Sarbecoviruses showed more amino-acid-changing mutations within SARS-CoV-2 (Spearman correlation 0.70), indicating Sarbecovirus evolutionary pressures still apply during the current pandemic (**Fig. 5a**). This inter-vs-within-strain agreement also held at codon resolution, with amino-acid-changing mutations preferentially disrupting non-conserved residues (535 mutations in 3264 positions, 16.4%) vs. conserved residues (607 in 6480, 9.4%, p<10^−10^) (**Extended Data Fig. 9a**).

### Accelerated and decelerated evolution

Notable deviations from this general agreement may reflect recent accelerated/decelerated evolution. S1 showed significantly-fewer mutations than expected from its extremely-high inter-strain rate (13% amino-acid-changing mutations observed vs. 17% expected, nominal p=0.0017, depletion: 28); additional SNVs (N=2696, May 9, 2020) further strengthened the statistical significance of this result (p=0.00033). Nsp3 also showed significantly fewer mutations than expected (10% vs. 15%, nominal p<10^−9^, depletion: 90) and Nucleocapsid significantly more (21% vs. 11%, nominal p<10^−8^, excess: 42).

The lower-than-expected number of mutations in S1/nsp3 might indicate recent mutation-rate or selective-pressure changes, possibly stemming from different phases of host-adaptive evolution, with pre-pandemic earlier-adapting S1/nsp3 (eg. via non-human-host transmission or undetected human transmission) requiring fewer pandemic-phase human-adaptive mutations than other later-adapting genes (noting that only a subset of mutations are adaptive). Alternatively, S1/nsp3 may have more positions in which deleterious mutations would be strongly-deleterious (purified-out even in shorter timescales) vs. mildly-deleterious (purified-out only over larger timescales). Lastly, frequent S1 recombination could inflate inter-strain rate estimates, but probably insufficiently to account for the observed discrepancies. (**Supplementary Text S6**).

The higher-than-expected number of variants in N might be explained by positive selection for host adaptation. We investigated whether such positively-selected variation might be clustered in specific segments, and searched the entire genome for clusters of variants disrupting conserved amino acid residues. We found no significantly-depleted regions and only one region significantly-enriched (**Supplementary Text S7**) relative to gene-specific variant density (p<0.012 after conservative genome-wide multiple-hypothesis correction), which was indeed localized in N, and contained 14 variants disrupting conserved residues (out of the observed excess of 29 such variants in N) concentrated in 20-amino-acid region R185-G204 (noting this enrichment is relative to the already-high enrichment of such variants in N). This region overlaps a predicted B-Cell epitope^43^, suggesting positive selection for immune system avoidance (**Fig. 5b, Extended Data Fig. 9c**).

### Spike SNV prioritization

We next investigated whether we can help prioritize candidate driver SNVs associated with phenotypic differences between SARS-CoV-2 strains, using the evolutionary history of each amino acid across Sarbecoviruses to provide position-specific estimates of evolutionary constraint, thus taking into account the biological context and precise functions that each amino acid plays in coronavirus biology (beyond position-independent general estimates from general amino acid properties).

As proof-of-principle, we focused on 16 amino-acid-changing variants in Spike with high frequency and/or epitope proximity^44,45^ (**Supplementary Table S3**). Among them, radical-amino-acid-change D614G, which rose in frequency across multiple cities and increases infectivity *in vitro*^*45–47*^, disrupts a perfectly-conserved residue (across Sarbecoviruses), and lies in a stretch of 11 perfectly-conserved amino acids (**Fig. 5c**), indicating its disruption is deleterious in bat-host contexts, and likely represents a novel human-host adaptation.

Of the other 15 Spike variants, two are in perfectly-conserved residues (V615I/F, P1263L) and two in mostly-conserved residues in highly-conserved regions (A831V, A829T/S), indicating likely-functional changes. Another three are in moderately-conserved contexts (V367F, D839Y/N/E, D936Y/H) less likely to be functional, and eight lie in repeatedly-altered amino acids in poorly-conserved regions and likely-neutral.

Lastly, Sarbecovirus evolutionary context helps prioritize likely drivers among co-inherited mutations. Spike D614G was nearly always co-inherited with Pol P4715L (also radical and altering a perfectly-conserved residue in a highly-conserved context, but potentially-deleterious given Pol’s slow evolution and less-likely-to-be-adaptive function), nsp3 nucleotide change C3037T (repeatedly-observed synonymous change, outside synonymously-constrained elements, likely-neutral), and nucleotide change C241T (perfectly-conserved, non-coding, in a loop of six unpaired bases in the conserved 5’-UTR SL5B secondary structure^34^ 25 nucleotides upstream of ORF1ab).

### Synonymous and non-coding substitutions

Even for synonymous SNVs we found agreement between cross-strain and within-strain constraint, with synonymously-constrained codons showing fewer synonymous variants (73 of 1394, 5.2%) than non-synonymously-constrained codons (555 of 8350 positions, 6.6%, binomial p=0.029, **Extended Data Fig. 9b**).

We also classified 643 intergenic and 5’/3’-UTR positions as “conserved” (N=432, 67%) or “non-conserved” (**Supplementary Table S3**), and found a surprising (but non-significant) SNV excess in conserved positions (17.4% vs. 13.7%, p=0.17).

## Discussion

We used comparative genomics to determine the conserved functional protein-coding genes of SARS-CoV-2, resulting in a new high-confidence evolutionarily- and experimentally-supported reference gene set, including ORFs 1a, 1ab, S, 3a, 3c, E, M, 6, 7a, 7b, 8, N, and 9b, but excluding 3b, 3d, 9c, and 10. We show that novel ORF 3c is functional and conserved, and that no other conserved genes remain to be discovered.

Our comparative genomics evidence complements experimental approaches by providing a comprehensive function-centric view of protein constraint, summed over all environmental conditions and hosts spanned by the strains compared here, while experimental methods only profile a single environmental and host condition in each experiment. Moreover, while experimental methods can suffer from incidental transcriptional or translational events, evolutionary signatures specifically measure functional constraint for a given function. While in principle our methods may miss recently-evolved genes that only function in a subset of strains, we found that our Sarbecovirus cross-strain evolutionary evidence agreed with SARS-CoV-2/SARS-CoV within-strain experimental evidence, suggesting it is unlikely that we may have missed newly-evolved genes.

It is important to note that comparative genomics methods that focus on nucleotide-level constraint such as phyloP and phastCons, as valuable as they are, would have mistakenly rejected S1 and ORF8 as seemingly non-conserved (given their extremely-rapid evolutionary rate and recombination history), and conversely included ORF10 as seemingly-conserved (given high nucleotide-level conservation in the overlapping RNA structure). Instead, our methods were able to correctly distinguish the protein-coding status of these genes because they use protein-coding evolutionary signatures that: (a) focus on the patterns of change characteristic of protein-coding constraint (specific codon substitution frequencies and reading frame conservation) rather than the overall number of substitutions; and (b) are less sensitive to the specific phylogenetic tree relating the genomes compared, and thus resilient to the frequent recombination events that characterize coronavirus genomes.

We found that both protein-coding and non-coding constraint agree between cross-strain Sarbecovirus substitutions and within-strain SARS-CoV-2 mutations, enabling us to classify SARS-CoV-2 variants into likely-functional vs. likely-neutral according to their evolutionary constraint. This revealed that the Spike D614G substitution likely represents a new adaptation to human hosts, as it disrupts a Sarbecovirus-conserved residue in a strongly-conserved region of S1, and to interpret the likely functional impact of genetic variants co-inherited with D614G based on their evolutionary history. Beyond the specific examples cited here, our annotations are broadly useful for interpreting SARS-CoV-2 variants and inferring causal relationships between viral mutations and disease phenotype. For interpreting future variants, we also created a genome browser track hub to facilitate SARS-CoV-2 variant interpretation based on their evolutionary context, and based on our revised gene annotations.

We found three notable exceptions to the otherwise-strong agreement between inter-strain and within-strain variation: N showed significantly more amino-acid-changing mutations than expected, and nsp3 and S1 showed significantly fewer. For N, the acceleration is consistent with positive selection for human-host adaptation across many variants, including a 20-amino-acid region enriched for conserved-residue-disrupting variants in a B-cell epitope. For nsp3 and S1, the deviation raises the possibility they may represent pioneer proteins that adapt to new-host transmission prior to its pandemic phase, then require fewer mutations while other proteins ‘catch up’, an observation that may be more generally true across different proteins showing acceleration/deceleration in different phases of host adaptation and pandemic spread. Another possibility is that the space of deleteriousness across all possible mutations is differently-distributed for nsp3 and S1 compared to other proteins, with more deleterious mutations in the strongly-deleterious end of the distribution, thus explaining the discrepancy in the number of observed amino-acid-changing substitutions between the short timescales captured in the recent pandemic SNVs vs. the longer timescales captured in cross-Sarbecoviruses comparative genomics. We discuss these and other possibilities in **Supplementary Text S6**.

Overall, our new reference gene set provides a solid foundation for systematically dissecting the function of SARS-CoV-2 proteins, and focusing experimental work on high-confidence uncharacterized ORFs, which can be guided in part by their evolutionary dynamics (such as the rapid evolution and recombination history of ORF6 and ORF8, indicating possible adaptive roles). In addition, our gene-level, codon-level, and nucleotide-level Sarbecovirus constraint, and the classification of all existing and potential SNVs into likely-functional vs. likely-neutral based on their evolutionary history, provide important foundations for elucidating SARS-CoV-2 biology, understanding it evolutionary dynamics, prioritizing candidate drivers mutations among co-inherited mutations, and prioritizing candidate regions for vaccine design and refinement.

## Supporting information

Summary Of Supplementary Data

CodAlignView PDFs For All Genes

Nextstrain Metadata

44 Sarbecovirus Alignment

44 Sarbecovirus Tree

PhyloCSF Genes.bed

PhyloCSF Rejected Genes.bed

## Methods

### Genomes and Alignments

Genome sequences were obtained from https://www.ncbi.nlm.nih.gov/. The genomes and NCBI annotations for SARS-CoV-2 and SARS-CoV were obtained from the records for accessions NC_045512.2 and NC_004718.3, respectively. The UniProt annotations for SARS-CoV-2 were obtained from the UCSC Genome Browser ^48^ on April 5, 2020.

The 44 Sarbecovirus genomes used in this study were selected starting from all betacoronavirus and unclassified coronavirus full genomes listed on ncbi via searches https://www.ncbi.nlm.nih.gov/nuccore/?term=txid694002[Organism:exp] and the same with txid1986197 and txid2664420 on 5-Mar-2020, excluding any that differed from NC_045512.2 in more than 10,000 positions in a pairwise alignment computed using NW-align ^49^, that cutoff being chosen so as to distinguish Sarbecovirus genomes among those that were classified, and removing near duplicates, including all SARS-CoV and SARS-CoV-2 genomes other than the reference. Coronavirus genomes in the left half of Extended Data Fig. 2 were those listed by https://www.ncbi.nlm.nih.gov/genomes/GenomesGroup.cgi?taxid=11118 on 11-Feb-2020.

The genomes were aligned using clustalo^50^ with the default parameters. The Phylogenetic tree was calculated using RAxML ^51^ using the GTRCATX model.

### PhyloCSF, FRESCo, and other conservation metrics

PhyloCSF (Phylogenetic Codon Substitution Frequencies)^22^ determines whether a given nucleotide sequence is likely to represent a functional, conserved protein-coding sequence by determining the likelihood ratio of its multi-species alignment under protein-coding and non-coding models of evolution that use pre-computed substitution frequencies for every possible pair of codons, and codon frequencies for every codon, trained on whole-genome data. PhyloCSF was run using the `29mammals` empirical codon matrices but with the Sarbecovirus tree substituted for the mammals tree. Input alignments were extracted from the whole-genome alignment and columns containing a gap in the reference sequence were removed. Browser tracks were created as described previously ^26^. Scores listed in Supplementary Table S2 were calculated on the local alignment for each ORF or mature protein, excluding the final stop codon, using the default PhyloCSF parameters, including `-- strategy=mle`.

FRESCo ^29^ was run using HYPHY version 2.220180618beta(MP) for Linux on x86_64 on 9-codon windows in each of the NCBI annotated ORFs. Alignments were extracted for the ORF excluding the final stop codon, and gaps in the reference sequence were removed. SCEs were found by taking all windows having synonymous rate less than 1 and nominal p-value<10^−5^, and combining overlapping or adjacent windows. For the variant analysis, FRESCo was also run on 1-codon windows using codon alignments as described previously ^29^.

Substitutions per site and per neutral site for each annotated ORF and mature protein were calculated by extracting the alignment column for each site or, respectively, 4-fold degenerate site, from the whole-genome alignment and determining the parsimonious number of substitutions using the whole-genome phylogenetic tree. For columns in which some genomes did not have an aligned nucleotide, the number of substitutions was scaled up by the branch length of the entire tree divided by the branch length of the tree of genomes having an aligned nucleotide in that column.

PhastCons and phyloP tracks shown in Fig. 2 are the Comparative Genomics tracks from the UCSC Genome Browser, which were constructed from a multiz ^52^ alignment of the list of 44 Sarbecovirus genomes that we supplied to UCSC.

### Analysis of Single Nucleotide Variants

Single nucleotide variants were downloaded from the “Nextstrain Vars” track in the UCSC Table Browser on 2020-04-18 at 11:46 AM EDT. Table S3 includes one additional mutation, G24047A, from a later download, in order to represent Korber variant A829T/S. We defined an amino acid to be “conserved” if there were no amino-acid-changing substitutions in the Sarbecovirus alignment of its codon. We defined codons to be “synonymously constrained” if the synonymous rate at that codon calculated by FRESCo using 1-codon windows was less than 1.0 with nominal p-value<0.034, corresponding to a false discovery rate of 0.125. We defined an intergenic nucleotide to be “conserved” if there were no substitutions of that nucleotide in the Sarbecovirus alignment. We classified SNVs as Synonymous, Nonsynonymous, or Noncoding, relative to the NCBI annotations, so SNVs within ORF10 were classified as coding, and SNVs within overlapping ORFs 3c and 9b were classified relative to the longer containing ORFs 3a and N, respectively. However, in Supplementary Table S3, we also classified variants according to our proposed reference gene annotations (fields beginning with New_); when classifying variants in overlapping ORFs 3a/3c and N/9b we classify SNVs relative to the ORF in which the variant is non-synonymous if that is true for only one of the frames, or the ORF for which the amino acid change is more radical (as defined by the blosum62 matrix obtained from biopython version 1.58 53) if it is non-synonymous in both frames, or the larger ORF if the variant is synonymous in both frames.

We determined mature proteins for which the density of amino-acid-changing SNVs differed significantly from the density that would be expected from their level of conservation, by calculating the residual of a linear regression of amino-acid-changing SNV density as a function of the fraction of conserved amino acids, for all mature proteins. The regression line was y=0.235-0.165x. We determined significance using a binomial p-value with a false discovery rate cutoff of 0.05. To further test significance of the SNV depletion in S1, we downloaded a larger set of SNVs from the UCSC Table Browser as above on 2020-05-09.

The 16 Spike-protein variants prioritized were those reported by Korber et al. in their bioRxiv preprint or later *Cell* publication (ones at greater than 0.3% frequency, or 0.1% if near certain epitopes).

To find regions that were significantly enriched for missense variants in conserved amino acids, we first defined a null model as follows. For each mature protein, we counted the number of missense variants and the number of conserved amino acids and randomly assigned each SNV to a conserved amino acid in the same mature protein, allowing multiplicity. For any positive integer n, we found the largest number of variants that had been assigned to any set of n consecutive conserved amino acids within the same mature protein across the whole genome. Doing this 100,000 times gave us a distribution of the number of missense variants in the most enriched set of n consecutive conserved amino acids in the genome. Comparing the number of actual missense variants in any particular set of n consecutive conserved amino acids to this distribution gave us a nominal p-value for that n. We applied this procedure for each n from 1 to 100 and multiplied the resulting p-values by a Bonferroni correction of 100 to calculate a corrected p-value for a particular region to be significantly enriched. We note that these 100 hypotheses are correlated because enriched regions of different lengths can overlap, so a Bonferroni correction is overly conservative and our reported p-value of 0.012 understates the level of statistical significance. To find significantly depleted regions we applied a similar procedure with every n from 1 to 1000, but did not find any depleted regions with nominal p-value less than 0.05 even without multiple hypothesis correction.

### Miscellaneous

Ribosome footprints shown in Extended Data Fig. 3 are from the track hub at ftp://ftp-igor.weizmann.ac.il/pub/hubSARSRibo.txt ^10^.

### Data Access

The PhyloCSF tracks and FRESCo synonymous constraint elements are available for the SARS-CoV-2/wuhCor1 assembly in the UCSC Genome Browser at http://genome.ucsc.edu as public track hubs ^1,23,24,48^ named “PhyloCSF” and “Synonymous Constraint”. The alignments and phylogenetic tree used here are included as supplementary materials. The alignments may be viewed, color coded to indicate protein-coding signatures, using CodAlignView (https://data.broadinstitute.org/compbio1/cav.php) with alignment set wuhCor1_c and chromosome name NC_045512v2.

Our proposed reference gene set for SARS-CoV-2 is included in BED format in Supplementary materials and is available as the “PhyloCSF Genes” track in the UCSC Genome Browser. A track showing the genes we have rejected may also be displayed using the configuration page.

A browser track showing SARS-CoV-2 single nucleotide variants, color coded by whether they are non-coding, synonymous, or amino-acid-changing, and whether they are in conserved codons, as well as tracks showing all codons that are conserved at the amino acid or synonymous level, may be viewed in the UCSC Genome Browser using the track hub at https://data.broadinstitute.org/compbio1/SARS-CoV-2conservation/trackHub/hub.txt. The details page for each SNV includes information about Sarbecovirus conservation and a link to view the alignment of a neighborhood of the SNV in CodAlignView. It is our intention to update this track hub as the list of variants in the UCSC Table Browser is updated. [Note to reviewers: classification is currently with respect to NCBI annotations; we will add a track classifying SNVs with respect to our PhyloCSF Genes annotations once our paper is accepted.]

In this resource, we have augmented variant data made available by UCSC ^54^ with our own annotations. UCSC data came from nextstrain.org ^55^, which was derived from genome sequences deposited in GISAID ^35^. Right of use and publication of the underlying sequences is entirely controlled by the authors of the original resource and the contributors of individual sequences, who are acknowledged in the Nextstrain metadata file included with supplementary materials. Our analysis provides an additional layer of annotation on their work rather than replicating or replacing it.

Original data usage policy as provided by UCSC: “The data presented here is intended to rapidly disseminate analysis of important pathogens. Unpublished data is included with permission of the data generators, and does not impact their right to publish. Please contact the respective authors (available via the Nextstrain metadata.tsv file) if you intend to carry out further research using their data. Derived data, such as phylogenies, can be downloaded from nextstrain.org (see “DOWNLOAD DATA” link at bottom of page) - please contact the relevant authors where appropriate.”

## Acknowledgements

We thank the UCSC genome browser staff and Maximilian Haeussler in particular for sharing our gene annotations with the community. We thank all contributors to the GISAID database for sharing primary sequences, and nextstrain.org/ucsc.edu for making variant data available. We thank Jeremy Luban, Robert Garry, and Mark Diekhans for helpful input. This work was supported by the National Human Genome Research Institute of the National Institutes of Health under Award Number U41HG007234. The content is solely the responsibility of the authors and does not necessarily represent the official views of the National Institutes of Health. Additional support was provided by the Wellcome Trust grant number WT108749/Z/15/Z and NIH grant R01 HG004037.

## Author Contributions

I.J. and M.K. conceived and designed the study and carried out all analyses. R.S. calculated synonymous constraint. I.J. and M.K. wrote the manuscript.

## Competing interest declaration

The authors declare no competing interests.

## Data Availability and Code Availability

All data generated or analysed during this study are included in this published article and its supplementary information files.

## Additional info

Supplementary Information is available for this paper.

Correspondence and requests for materials should be addressed to Manolis Kellis.

## Extended Data Figures

**Extended Data Figure 1.**
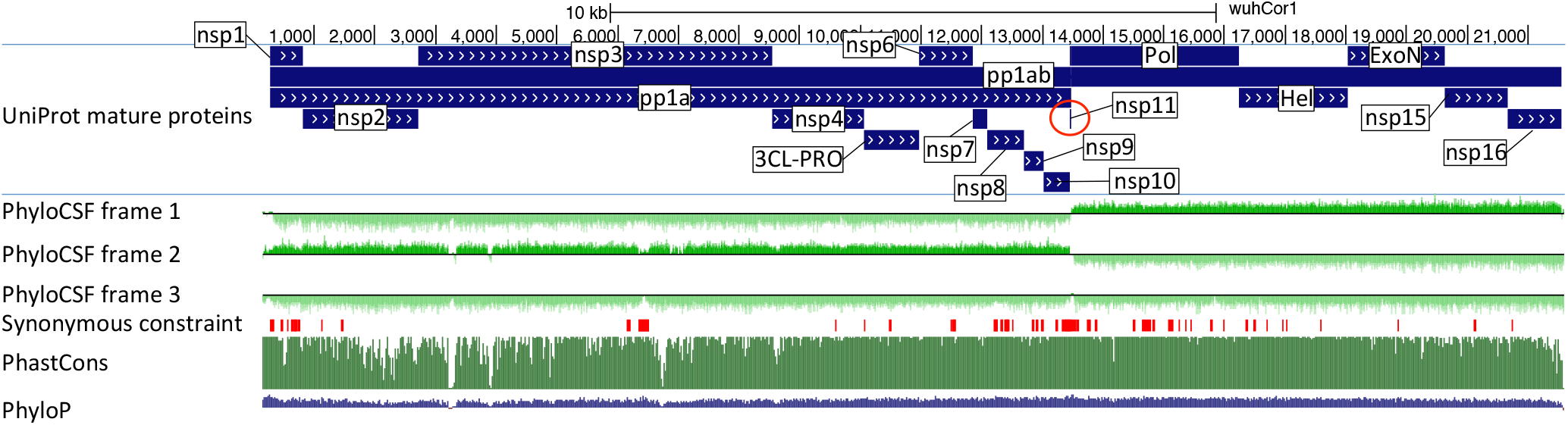
PhyloCSF signal for polyprotein ORF1. UCSC Genome Browser view SARS-CoV-2 genome for polyprotein ORF1ab showing UniProt gene annotations for individual non-structural proteins (nsp), PhyloCSF tracks (green) in each of 3 reading frames, and Synonymous Constraint Elements (SCEs, red), along with phastCons/phyloP nucleotide-level constraint (green/blue). Polyprotein 1ab is processed into 16 mature peptides nsp1-nsp16. PhyloCSF signal shows clear protein-coding signal for all proteins, indicating clearly that all are functional proteins (except nsp11, red circle, discussed in the main text). PhyloCSF signal captures the correct frame throughout the entire length of each protein (except nsp3, where some small regions show reduced frame-2 signal and/or increased frame-3 signal, but upon inspection these are only stop-codon-free in frame-2 and do not represent dual-coding candidates).

**Extended Data Figure 2.**
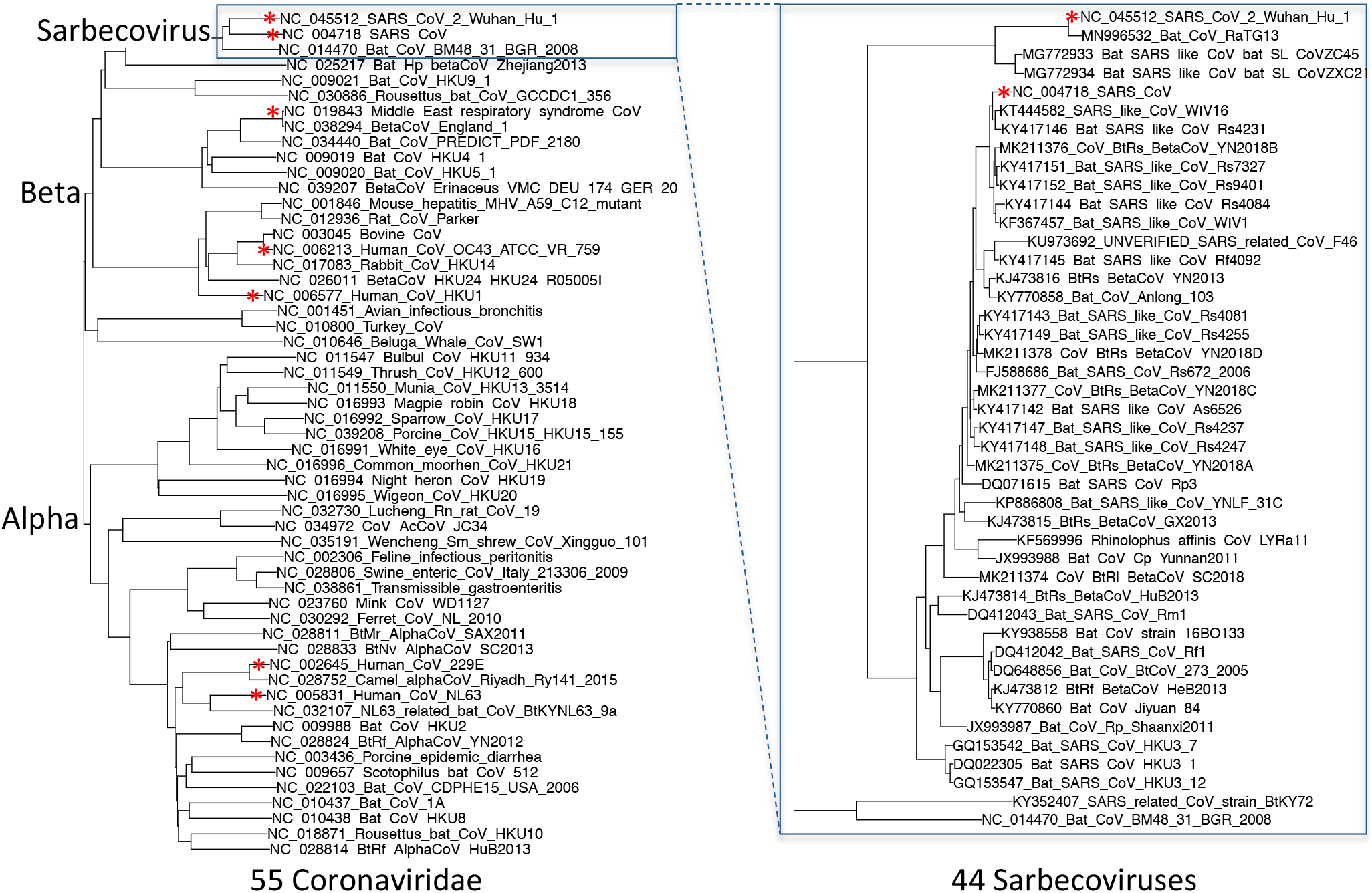
Phylogenetic tree of 44 Sarbecovirus genomes and larger phylogenetic context. Left: Phylogenetic tree of a selection of Coronaviridae genomes, including the seven that infect humans (red asterisks). Right: Phylogenetic tree of the 44 Sarbecovirus genomes used in this study. Trees are based on whole-genome alignments and might be different from the history at particular loci, due to recombination.

**Extended Data Figure 3.**
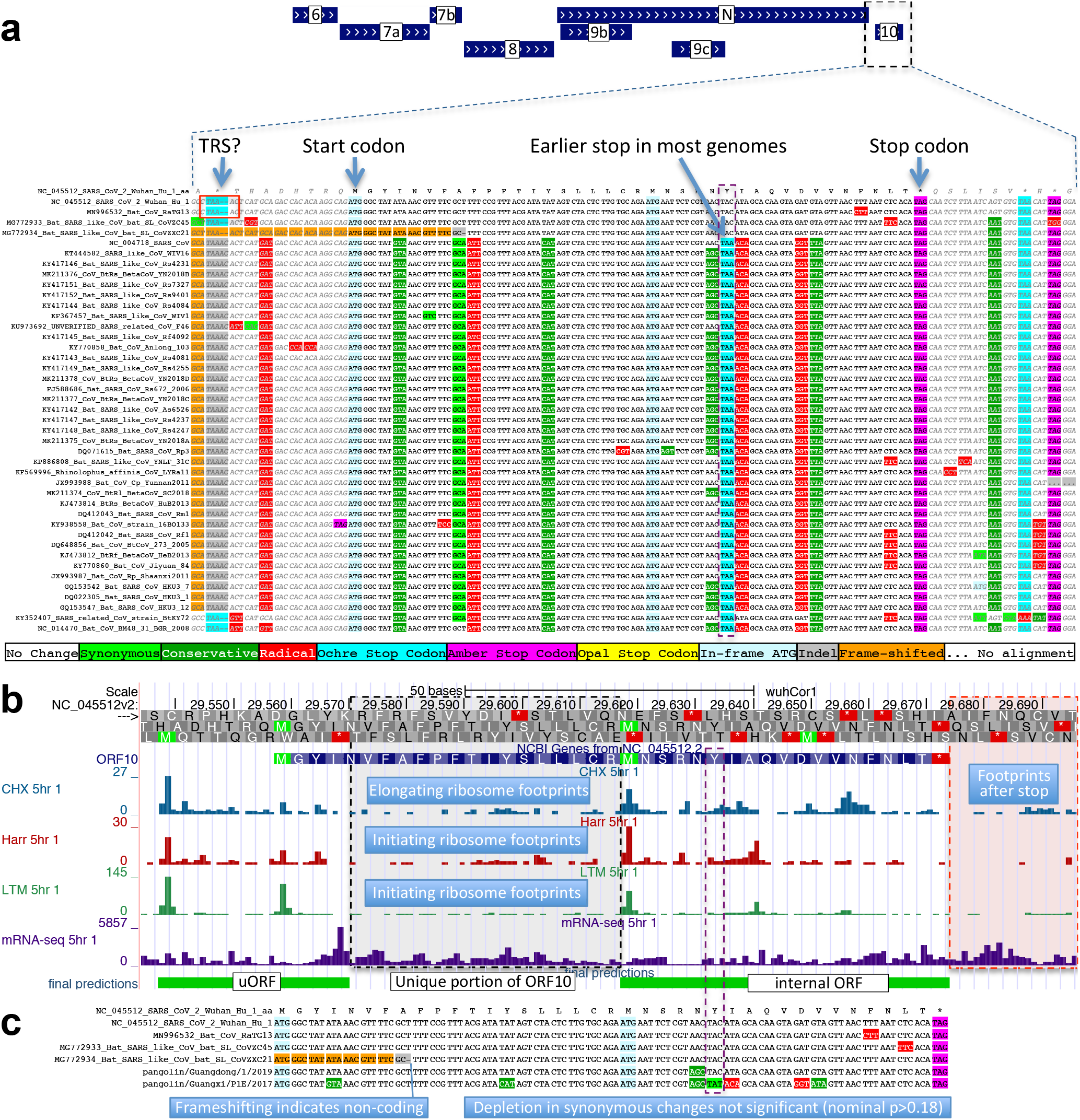
ORF10 is not protein-coding. **a.** Alignment of Sarbecovirus genomes at ORF10, including 30 additional flanking nucleotides on each side. Most substitutions are amino-acid-changing, either radical (red) or conservative (dark green), with only two synonymously-changing positions (light green), indicating this is not a protein-coding region. In addition, nearly all strains show an earlier stop codon (cyan), further reducing the length of this already-short ORF from 38 amino-acids to 25, and one of the four strains lacking the earlier stop includes a frame-shifting deletion. The putative partial transcription-regulatory sequence (TRS) present in SARS-CoV-2 and its closest relative (Bat CoV RaTG13) is not conserved in any other strains. The region surrounding ORF10 shows very high nucleotide-level conservation, which spans ORF10 and extends beyond its boundaries in both directions, indicating that this portion of the genome is functionally important even though it does not code for protein (indeed, this region is part of a pseudoknot RNA structure involved in RNA synthesis). **b.** Ribosome footprints previously used to suggest ORF10 translation^10^ in fact localize either in an upstream ORF (uORF, green) or in an internal ORF (green, “final predictions” track^10^), but not in the unique portion of ORF10 (dashed black box), indicating they are less likely to reflect functional translation of ORF10, and more likely to represent incidental translation initiation events. We note that the density of elongating footprints in the unique portion (black box) is no greater than the density after the stop codon (red box), consistent with incidental events. We also note that the internal ORF is only 18 codons long in 4 strains, and only 5 codons long in the other 40 Sarbecovirus strains, given the early stop codon (purple box) and unlikely to be functional. Footprint tracks show elongating ribosome footprints in cells treated with cycloheximide (blue, CHX), and footprints enriched for initiating ribosomes using harringtonine (Harr, red), and lactimidomycin (LTM, green). “mRNA-seq” track shows RNA-seq reads. **c.** CodAlignView^25^ of alignment previously used to argue that a high dN/dS ratio in ORF10 indicated positive selection for protein-coding-like rapid evolution^8^, based on only six closely-related strains (SARS-CoV-2, three bat viruses, two pangolin viruses). The authors noted a frameshifting deletion (orange/grey) in one of the bat viruses, which provides strong evidence against conserved protein-coding function, but they interpreted it (without evidence) as a potential sequencing error and excluded the strain from consideration. Even ignoring the frameshift-containing strain, the evidence used is insufficient to reach statistical significance: the alignment includes only 9 substitutions, of which 4 are radical, 4 are conservative, and 1 is synonymous. In a neutrally-evolving region with 9 substitutions, we would expect 2-3 synonymous changes, depending on the evolutionary model used, and a depletion to only 1 synonymous change is not statistically significant (nominal p-value>0.18 even in the most generous evolutionary model). This already-non-significant nominal p-value would move even further from significance with the necessary multiple-hypothesis corrections.

**Extended Data Figure 4.**
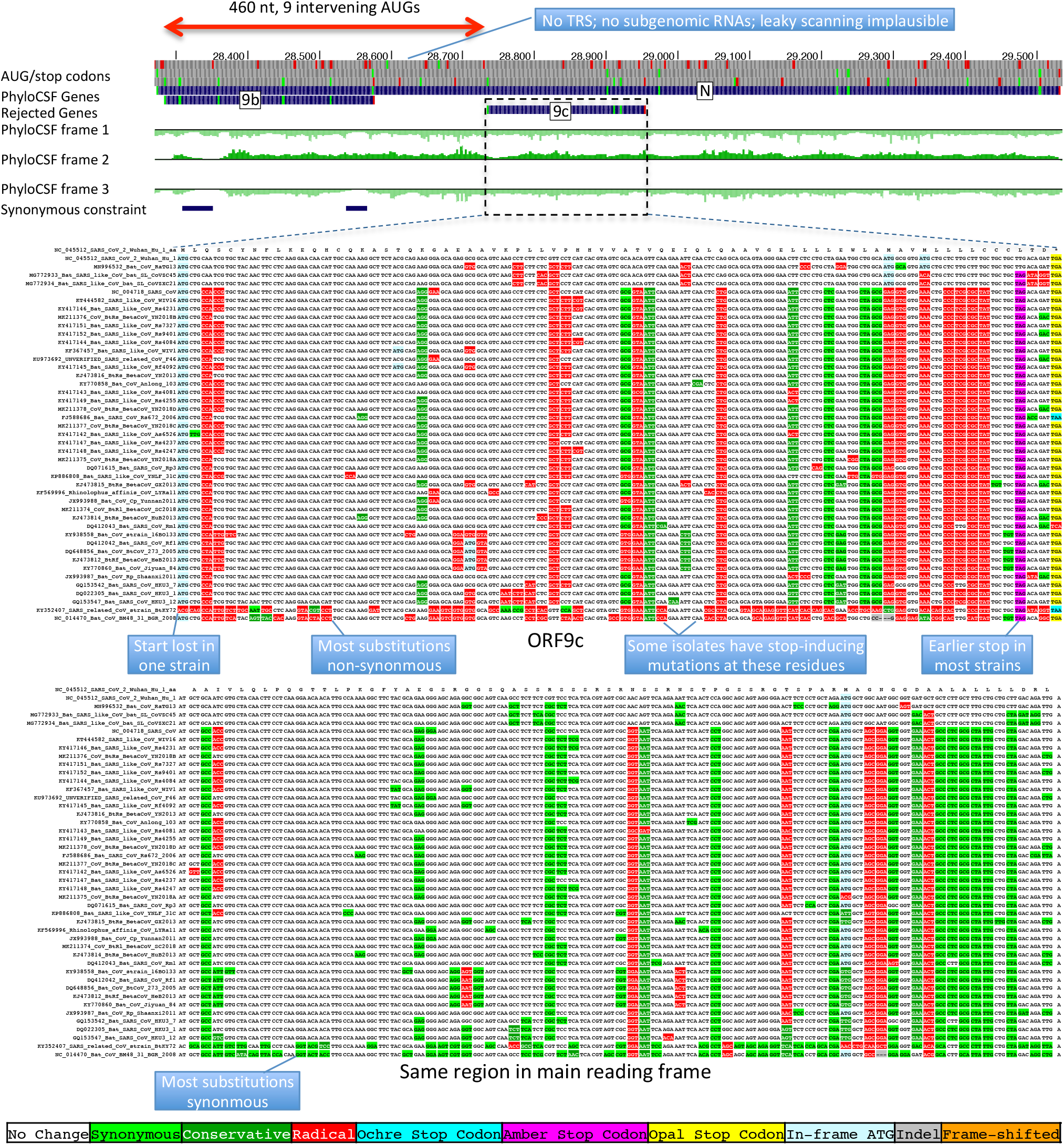
Nucleocapsid-overlapping ORF9c is not protein-coding. Sarbecovirus alignment of frame2-encoded ORF9c (top), which overlaps frame3-encoded Nucleocapsid (bottom). ORF9c start codon is lost in one strain, and most strains have an earlier UAG stop codon (magenta) 3 codons before the end. In Nucleocapsid-encoding frame 2 (bottom), nearly all nucleotide substitutions are amino-acid-preserving (synonymous, light green), indicating strong purifying selection for protein-coding function. By contrast, in ORF9c-encoding frame 3 (top), nearly all nucleotide substitutions result in function-disrupting (radical) amino acid changes (red), and very few result in synonymous (light green) or function-preserving (conservative, dark green) substitutions, indicating lack of purifying selection for protein-coding function for ORF9c, so it does not play conserved protein-coding functions. In addition, ORF9c is unlikely to be translated via leaky ribosome scanning because its start codon is 460 nucleotides after N’s (red arrow) with 9 intervening AUG codons (green dots), direct-RNA sequencing found no ORF9c-specific subgenomic RNAs^16–18^, no TRS is appropriately positioned to create one, and several SARS-CoV-2 isolates^35^ contain stop-introducing mutations^7^, indicating that ORF9c is not a recently-evolved strain-specific gene either. We conclude 9c is not protein-coding.

**Extended Data Figure 5.**
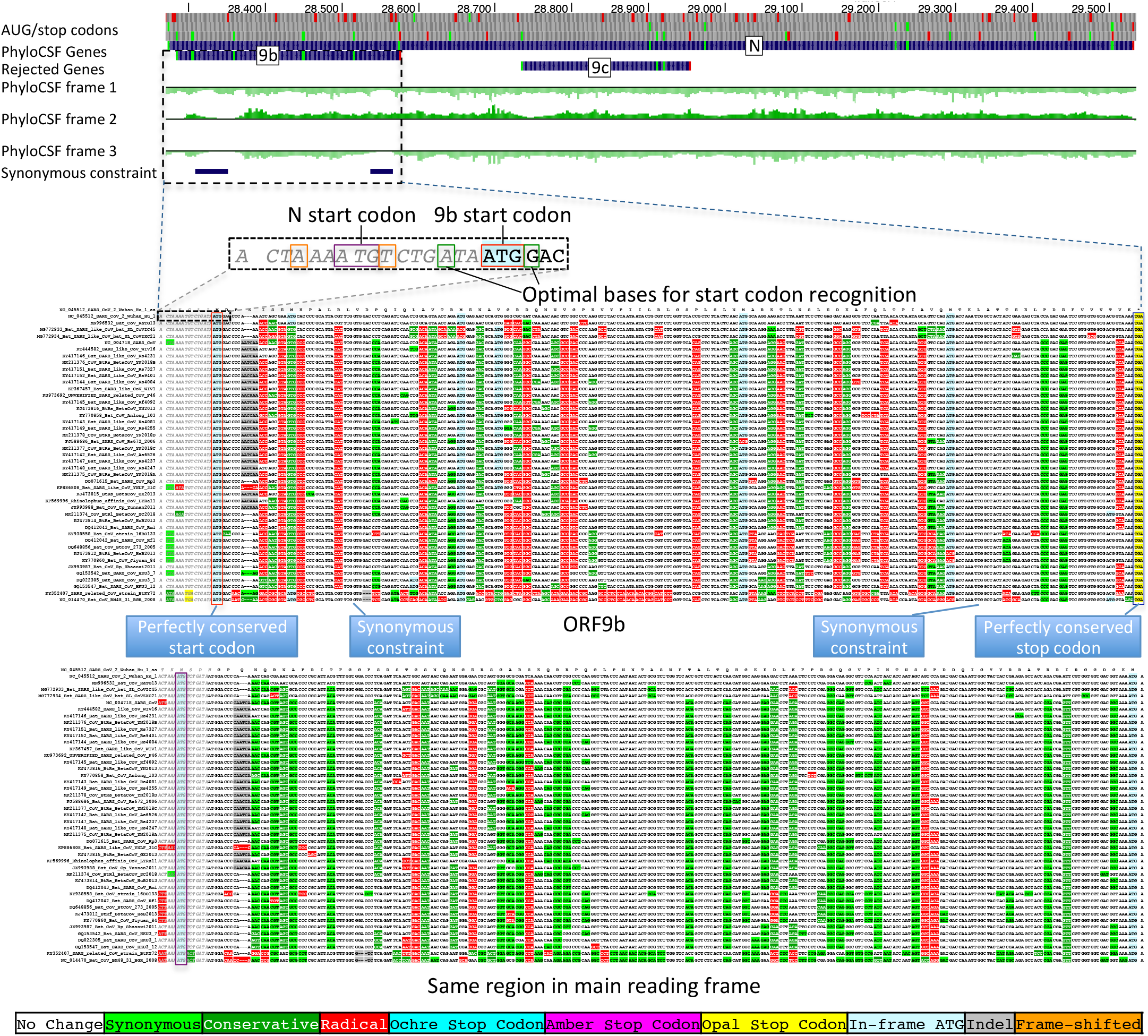
Nucleocapsid-overlapping ORF9b is protein-coding. Sarbecovirus alignment of frame3-encoded ORF9b (top), which overlaps frame2-encoded Nucleocapsid (bottom). Although ORF9b-encoding frame3 shows many function-disrupting (radical, red) substitutions, its start codon (red box) is perfectly conserved, its stop codon (blue box) is perfectly conserved, and there are no intermediate stop codons in any strain. Moreover, its Kozak start-codon context (dashed black box) is optimal for ribosomal start codon recognition, with A in position -3 and G in position +4 (green boxes), while the start codon context of N is less optimal, with an A in -3 and T in +4 (orange boxes), making it likely that ORF9b can be translated by leaky scanning from the same subgenomic RNA as N, as it is only ∼2 codons downstream of N’s start. Moreover, both the optimal 9b start-codon context, and the less-optimal N start-codon context are fully-conserved features across all Sarbecovirus strains, indicating that leaky-scanning translation may be a conserved feature throughout Sarbecoviruses. In addition, ORF9b shows significant localized synonymous constraint in N in its start and end regions (**Fig. 3**), even relative to the overall low synonymous rate of N, consistent with dual-coding functions. ORF9b also has proteomics support^15,36,37^ in SARS-CoV-2, including evidence of viral-RNA binding^38^, and alternate-frame translation support by ribosome profiling^10^. In SARS-CoV, ORF9b protein (and antibodies to it) was detected in SARS patients^39,40^, localized in mitochondria, and interfered with host cell antiviral response when overexpressed^41^. We conclude ORF9b encodes a conserved functional protein-coding gene.

**Extended Data Figure 6.**
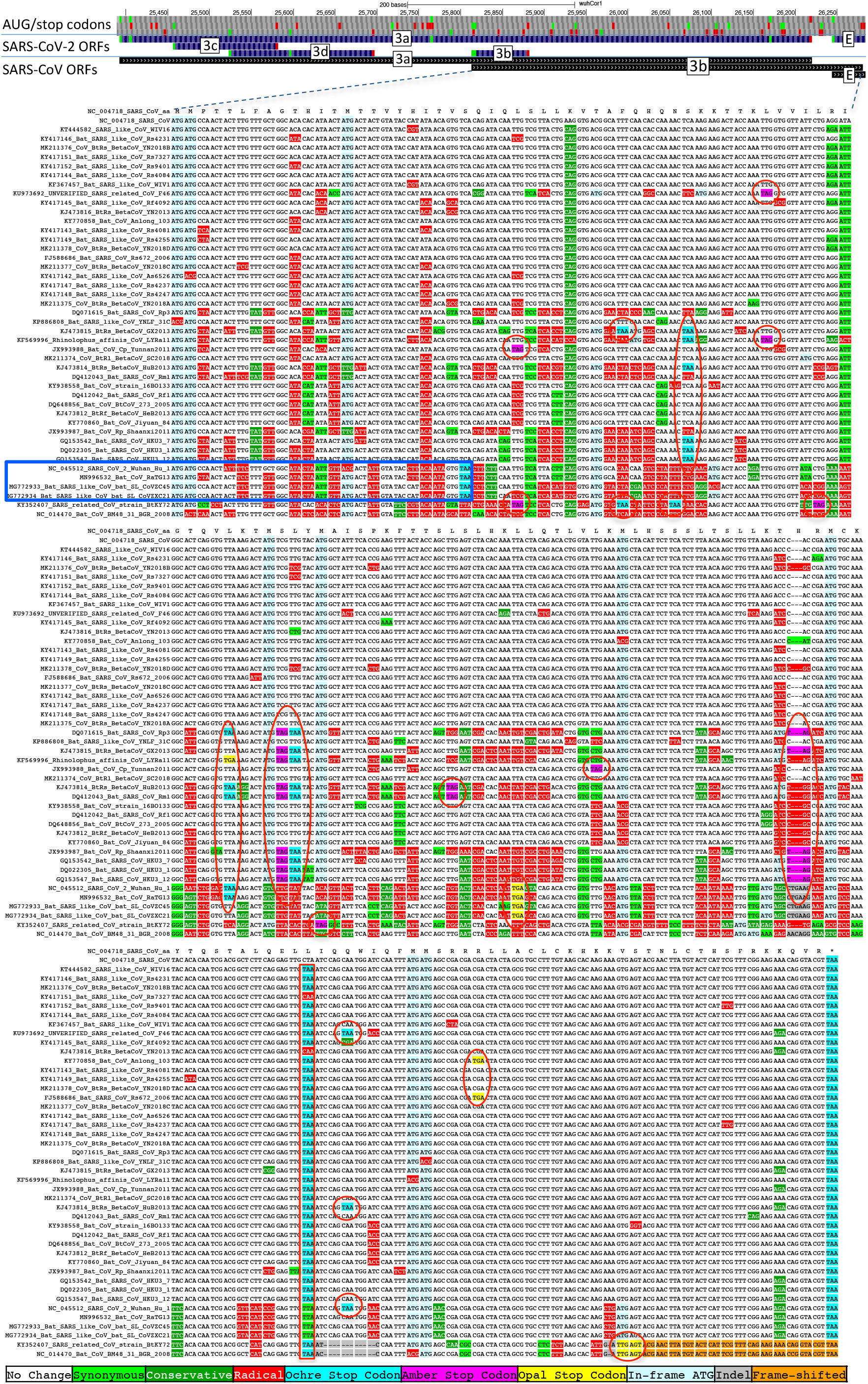
ORF3b is not protein-coding. Sarbecoviruses alignment of SARS-CoV 154-codon ORF3b overlapping ORF3a, (reordered with SARS-CoV and related strains on top). Although start codon is conserved in all but one strain, ORF length is highly variable due to numerous in-frame stop codons (red ovals and red rectangle). The 22-codon ORF in SARS-CoV-2 has strongly negative PhyloCSF score, does not overlap any SCEs, and even among the four strains sharing its stop codon (blue rectangle) all six substitutions are radical amino acid changes, providing no evidence of amino-acid-level purifying selection. Ribosome profiling did not find translation of ORF3b, transcription studies did not find substantial transcription of an ORF3b-specific subgenomic RNA, and translation by leaky scanning would implausibly require ribosomal bypass of eight AUG codons (green rectangles, top panel), some with strong Kozak context. (Supplementary Fig. S3 has comparison to reading frame of ORF3a.)

**Extended Data Figure 7.**
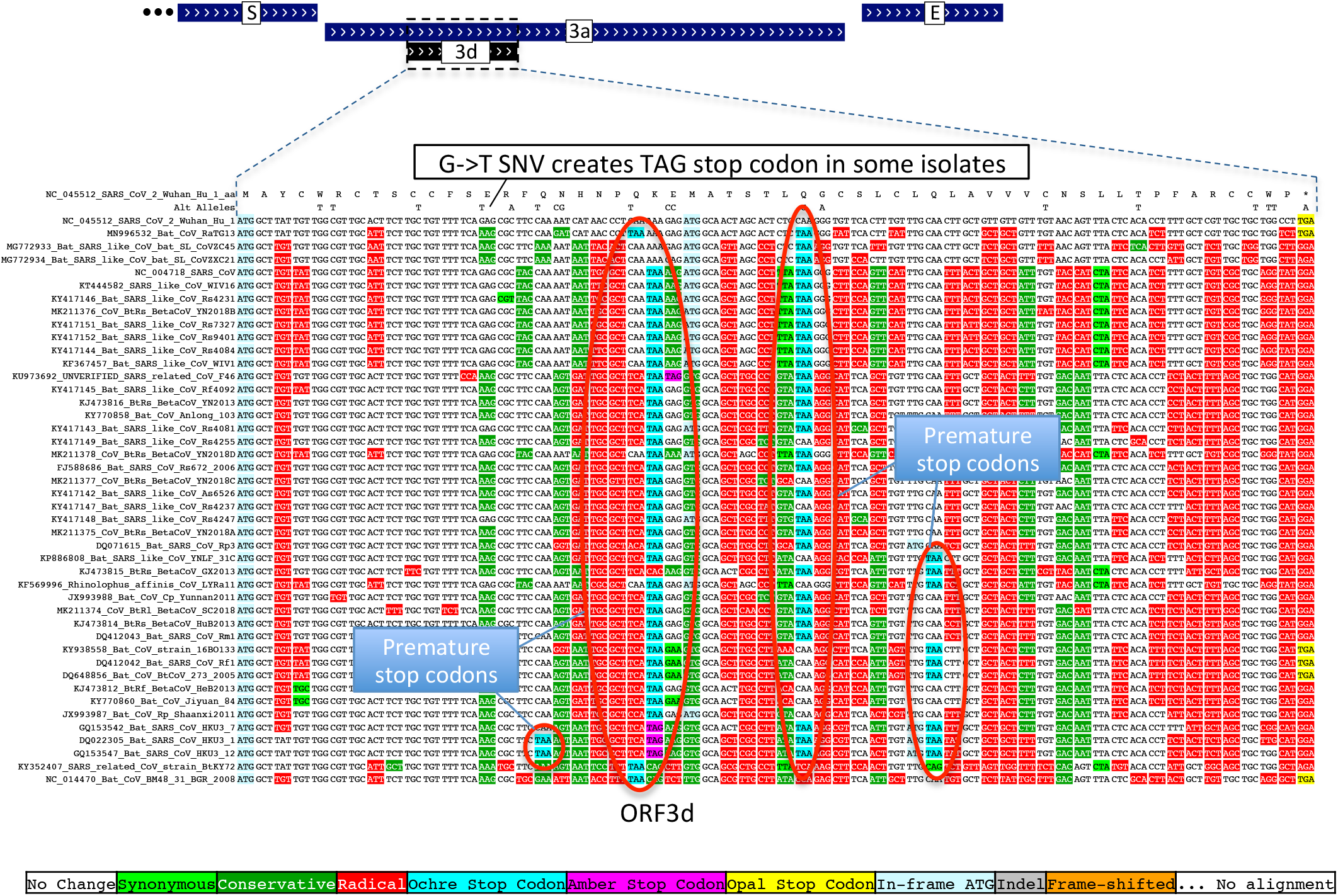
ORF3d is not protein-coding. Sarbecovirus alignment of 57-codon ORF3d (referred to by some authors as 3b) overlapping ORF3a shows mostly function-altering radical amino-acid substitutions (red columns), and repeated interruption of by one or more premature stop codons in all other strains (red ovals), unambiguously indicating that ORF3d is not a conserved protein-coding gene. A substantial fraction of SARS-CoV-2 isolates have stop-introducing mutations, and ribosome profiling did not identify ORF3d as a translated ORF^10^, indicating that it is not a recently-evolved strain-specific gene either.

**Extended Data Figure 8.**
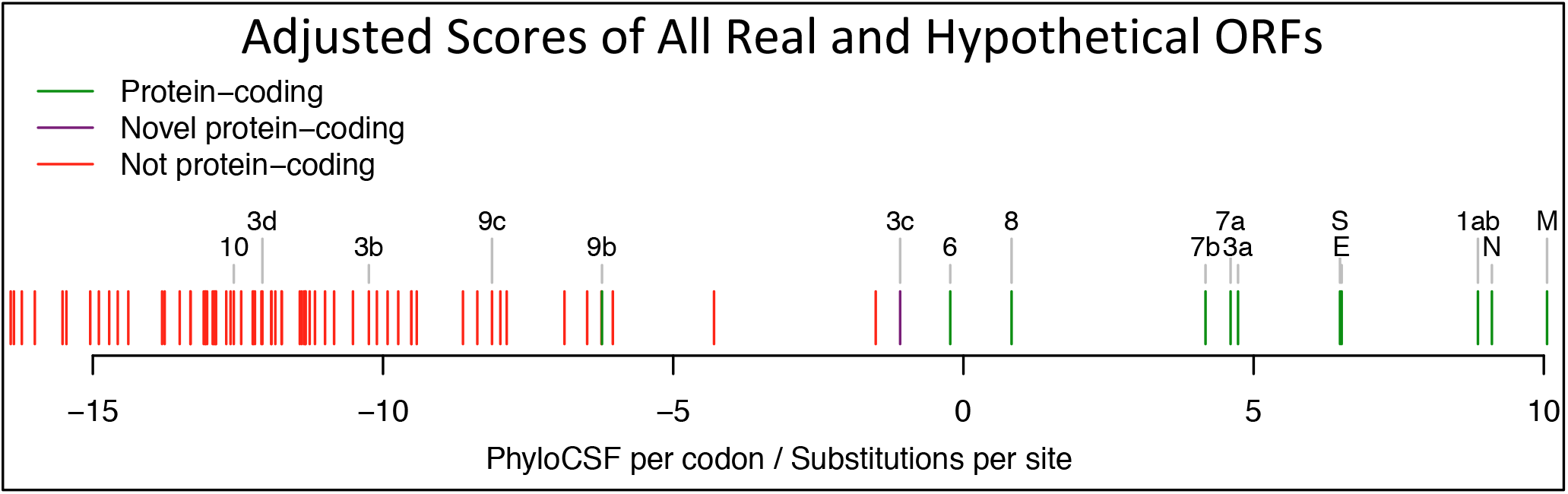
Branch-length-adjusted PhyloCSF score strongly rejects ORF10. Similar to Fig. 1c, but showing PhyloCSF scores per codon divided by the average number of substitutions per site, to adjust for the fact that high-nucleotide-conservation regions show compressed unscaled PhyloCSF scores (closer to zero) because there are fewer nucleotide substitution events. The branch-length-scaled score distribution further separates the scores of confirmed protein-coding genes (green) from non-protein-coding segments (red). The very low score of ORF10 with this adjustment indicates that its only-slightly-negative unscaled-PhyloCSF score in Fig. 1c stems from the high nucleotide conservation of the region, rather than protein-coding constraint. The scores of N-overlapping ORFs 9b and 9c are both reduced, consistent with the high nucleotide conservation of N. Notably, the branch-length-adjusted score for 3c remains high, consistent with its protein-coding nature, and despite the higher overall nucleotide conservation of its dual-coding region. We have manually inspected all other candidates with adjusted scores higher than 9c, and all are rejected (as not protein-coding): two are discussed in Supplementary Figure S4 (and are not protein-coding), and the remaining all show internal stop codons (and are not protein-coding).

**Extended Data Figure 9.**
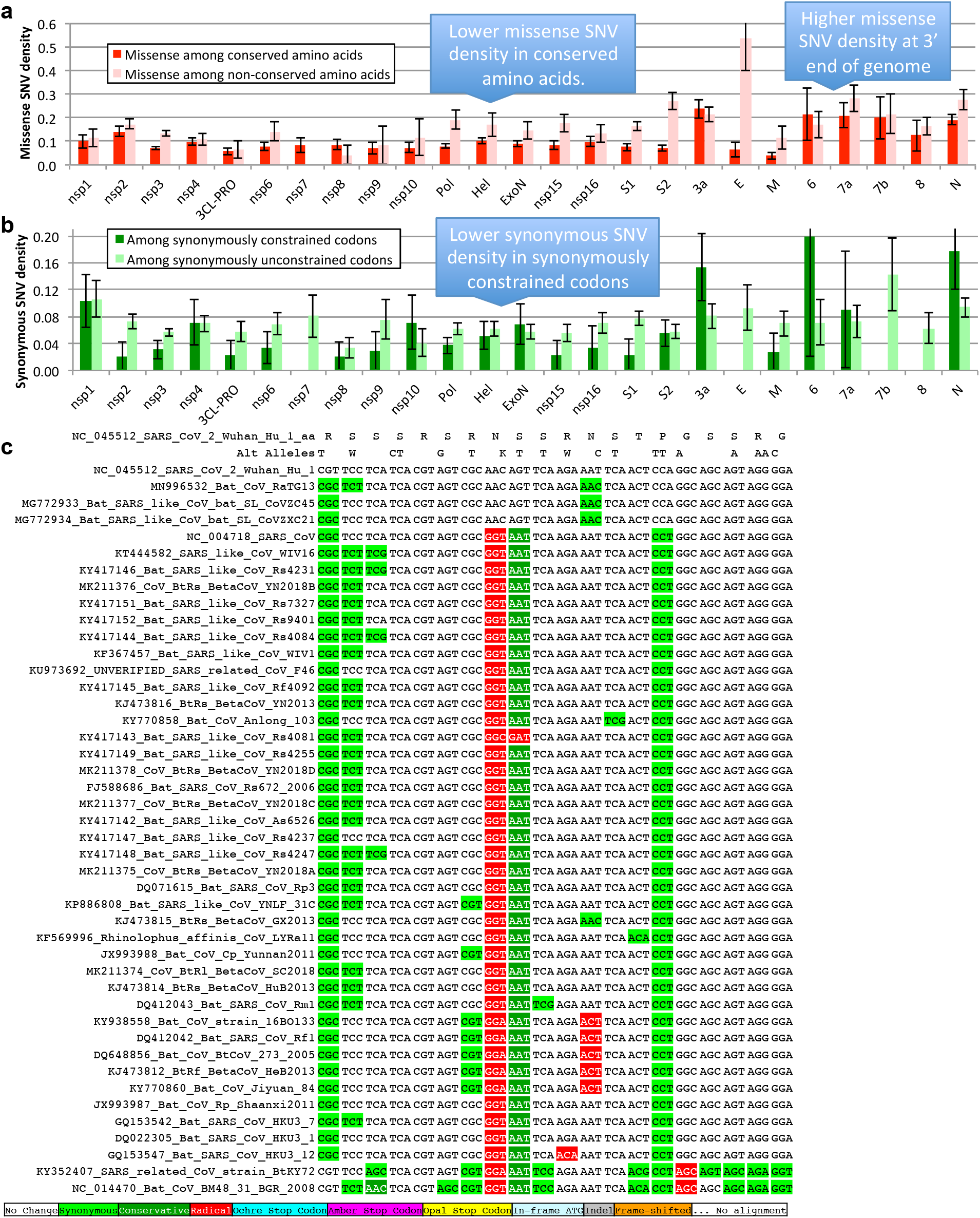
Single nucleotide variants and conservation. Error bars indicate standard error of mean. **a.** Density of SNVs disrupting conserved amino acids (dark red) is significantly lower than disrupting non-conserved amino acids (light red). Both densities are higher near the 3’ end of the genome, indicating higher mutation rate or less purifying selection even among amino acids that are perfectly conserved in Sarbecovirus. **b.** Density of synonymous variants in synonymously constrained codons (dark green) is significantly lower than among synonymously unconstrained codons (light green), a depletion seen in most genes. Overall, conservation in the Sarbecovirus clade at both the amino acid level and nucleotide level is associated with purifying selection on variants in the SARS-CoV-2 population. **c.** Alignment of 20 amino acid Nucleocapsid region that is highly enriched for variants disrupting perfectly conserved amino acids (alternate alleles shown in second row, W = A or T, K = G or T). There are 14 non-synonymous variants among the 14 perfectly conserved amino acids (columns with no red or dark green). This region is contained within a predicted B Cell epitope, suggesting positive selection for immune system avoidance.

## Notes

### Competing Interest Statement

The authors have declared no competing interest.

### Summary of Updates

- Proposed a new reference gene set for SARS-CoV-2. - Concluded that ORF9b is protein-coding. - Analyzed and rejected recently-proposed ORF3d and (SARS-CoV-2) ORF3b as not protein-coding. - Found that cross-strain and within-strain evolutionary pressures largely agree at the gene level with some notable exceptions, including fewer-than-expected mutations in nsp3 and Spike subunit S1, and more-than-expected mutations in Nucleocapsid. - Found that Nucleocapsid region enriched for missense mutations lies in a predicted B-cell epitope. - Renamed ORF14 -> ORF9c. - Removed analysis of coding potential of SARS-CoV genes not in SARS-CoV-2. - Removed analysis of variants from Yao et al predicted to affect viral load in-vitro. - Shortened and streamlined presentation.

