## Supplementary Figures S1-5 and Text S1-7 for "SARS-CoV-2 gene content and COVID-19 mutation impact by comparing 44 Sarbecovirus genomes"

Supplementary Text S6 Explanations for within-/cross-strains deviation in nsp3/S1 ..10

Supplementary Text S7 Enriched and depleted clusters of missense SNVs..............11

### Supplementary Figures

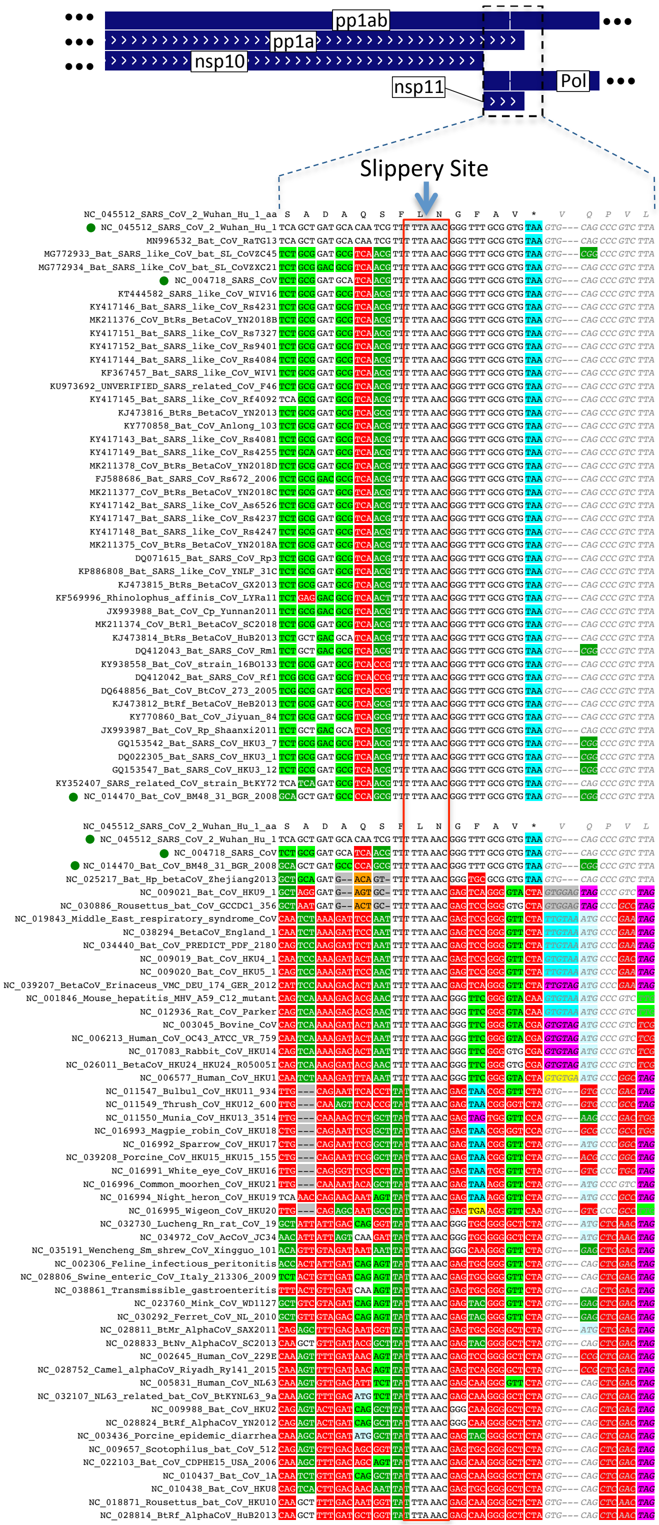

**Supplementary Figure S1. Alignment of nsp11 and frameshift site.**

Alignment of nonstructural protein nsp11 and subsequent 5 codons in 44 Sarbecoviruses (top) and 52 coronaviridae (bottom, green dots for strains included in both). Programmed frameshift slippery site (red rectangle) is perfectly conserved in all genomes. Pol shares the 5' nine codons of nsp11 but then continues 3' of the slippery site in a different reading frame. The four codons 3' of the slippery site are perfectly conserved in Sarbecoviruses, consistent with a dual coding region, but nsp11 is only 13 codons long and its stop codon is poorly conserved in coronaviridae (cyan, magenta, and yellow stop codons), suggesting that it does not produce a functional protein.

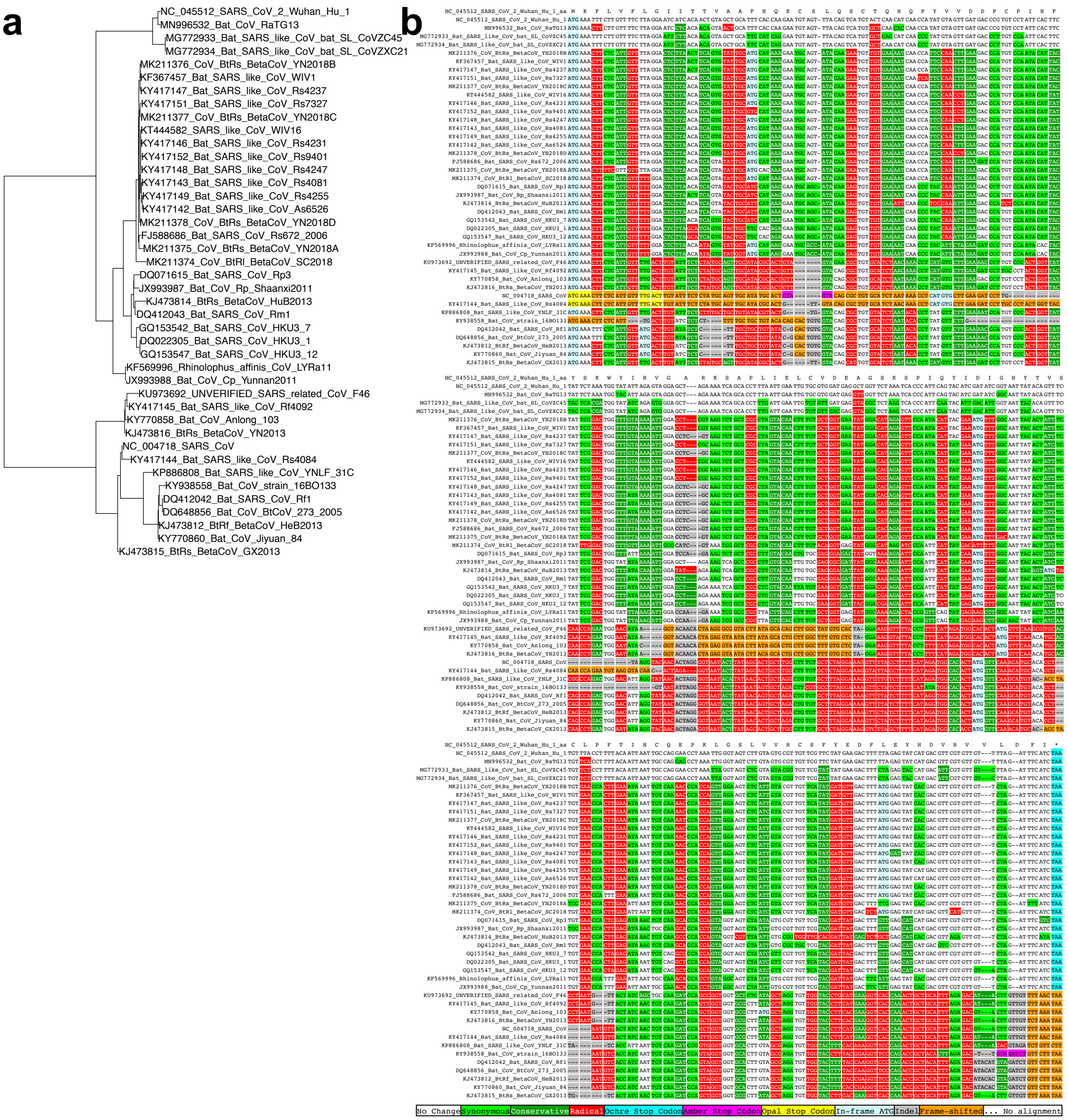

### Supplementary Figure S2. ORF8 Phylogeny.

**a.** Phylogenetic tree computed from ORF8 alignment differs from genome-wide tree in Extended Data Fig. 2, indicating there has been recombination. **b.** Alignment of ORF8 with strains ordered according to the ORF8 tree shows radical amino acid substitutions (red) disrupting almost every residue. Although the apparent high rate of nucleotide-level evolution in ORF8 when estimated using the whole genome tree (3.9 substitutions per site, 6.2 per four-fold degenerate site) is partly an artifact of estimation along a tree topology that does not match the history of ORF8, it remains very high (2.1 and 3.7 respectively, Supplementary Table S2) when estimated using the ORF8-specific phylogeny, substantially higher than other ORFs (e.g. 1.1 and 2.8 respectively for ORF1ab when restricting to the 42 strains aligned in ORF8), and thus is not entirely an artifact of recombination.

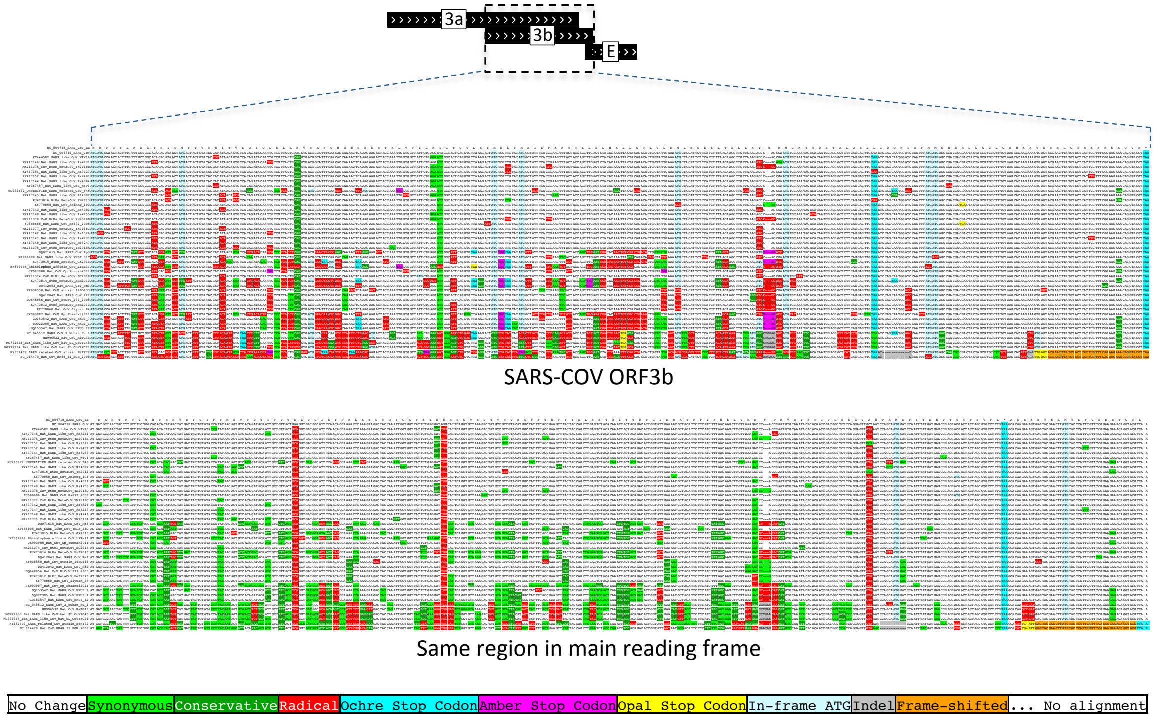

**Supplementary Figure S3. ORF3b alignment without wrapping.**
Alignment of SARS-CoV ORF3b (as in Extended Data Fig. 6 but without wrapping), and the same region in the reading frame of ORF3a.

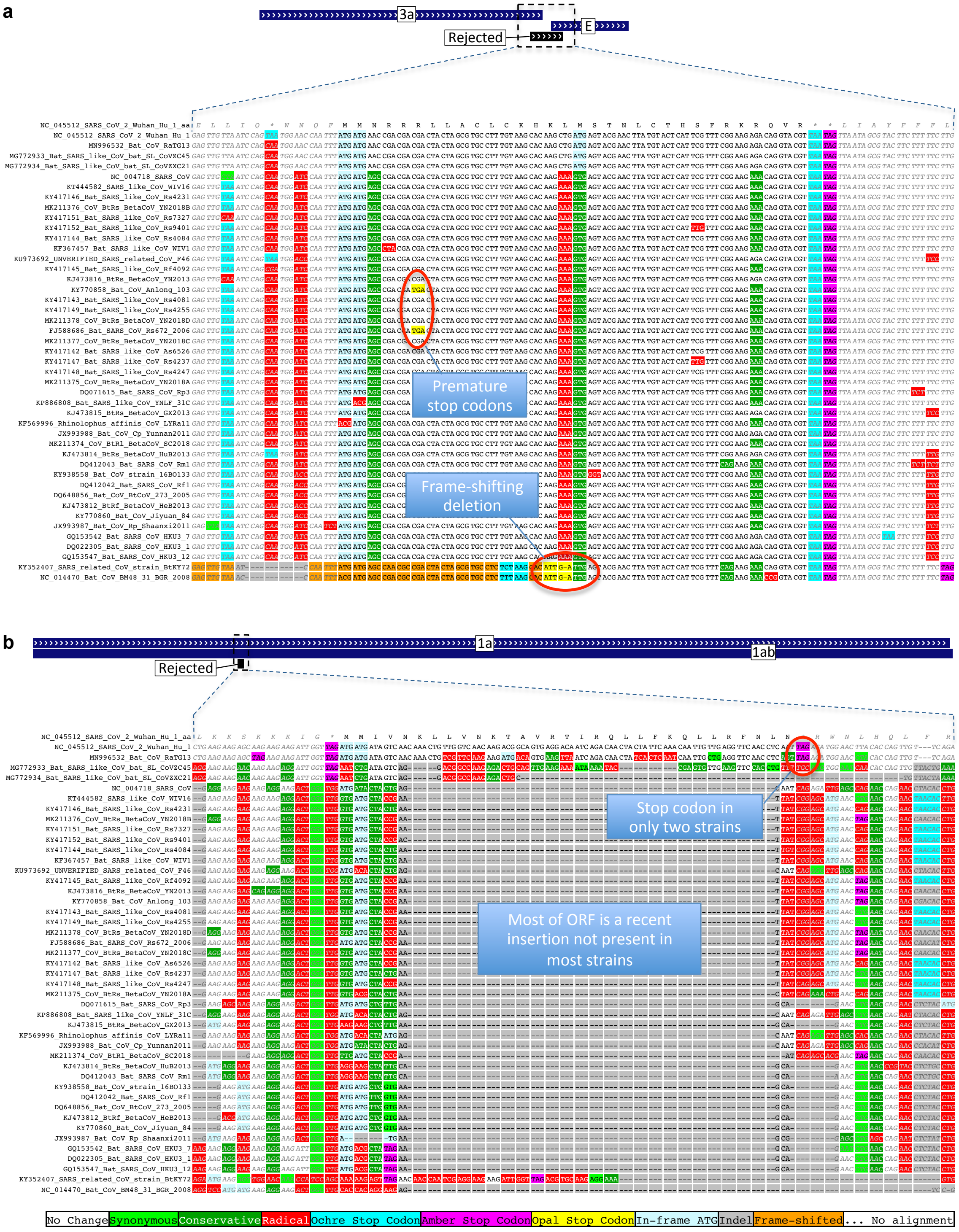

### Supplementary Figure S4. Alignments of rejected ORFs.

CodAlignView images of ORFs rejected during our search for novel conserved coding regions. **a.** 32-codon ORF (26183-26278) that overlaps the 3' end of ORF3a and the 5' end of E with PhyloCSF score -2.74. Two strains have a frame-shifting one-base deletion within the ORF, and two others have premature stop codons. None of the substitutions are synonymous. There is high nucleotide-level constraint, but it continues on both sides of the ORF, suggesting it does not result from translation of the ORF. This ORF is the 3' end of the SARS-CoV ORF3b orthologous region (which is interrupted by multiple stop codons in SARS-CoV-2). **b.** 31-codon ORF (3207-3299) overlapping ORF1a, with PhyloCSF score -7.77. Most of the ORF consists of a 75-nt insertion that is only present in SARS-CoV-2, RaTG13, and CoVZC45, and the start and stop codons are missing in CoVZC45, so this is not a conserved coding sequence.

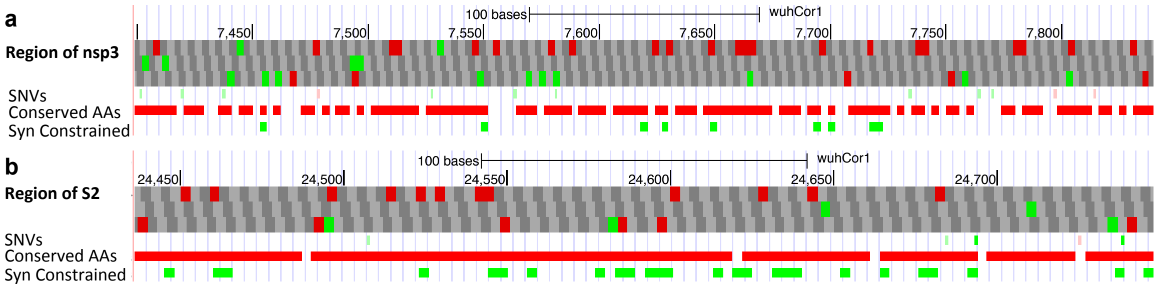

**Supplementary Figure S5. SNV-depleted regions.**
UCSC Genome Browser images of regions in nsp3 (**a)** and S2 (**b)**. Most amino acids in these regions are conserved (red rectangles in Conserved AAs track), but the only missense variants (light red rectangles in SNVs track) disrupt non-conserved amino acids (variants disrupting conserved amino acids would be bright red if present). The lack of missense variants in such a large set of conserved amino acid residues could indicate that constraint in the Sarbecovirus clade has continued particularly strongly in the SARS-CoV-2 population. However, although these are the most depleted regions in the genome for missense variants in conserved amino acid residues, neither depletion is statistically significant, even without any correction for multiple region lengths searched (nominal p=0.072 and p=0.093, respectively).

### Supplementary Text

#### Supplementary Text S1 Resolution of ambiguous gene names

The ORFs overlapping ORF3a and N have been referred to by different names by different authors, and different ORFs have been referred to by the same name. We have resolved inconsistently used names in communication with several other authors and with the approval of members of the ICTV Coronavirus Study Group. The ORFs, in 5' to 3' order, with our name, length, coordinates, and other names that have been used:

| Our name | Length (amino acids) | Coordinates (w/o stop) | Other names/Notes |
|---|---|---|---|
| ORF3c | 41 | 25457-25579 | ORF3h, 3a.iORF1 |
| ORF3d | 57 | 25524-25694 | ORF3b |
| ORF3b | 22 | 25814-25879 | 5’ end of SARS-CoV ORF3b ortholog |
| ORF9b | 97 | 28284-28574 | ORF9a, N.iORF1 |
| ORF9c | 73 | 28734-28952 | ORF9b, ORF14 |

#### Supplementary Text S2 Evidence that ORFs 3d and 3b are not protein-coding

ORF3b is a 22 amino acid ORF having coordinates 25814-25882, orthologous to the 5’ end of SARS-CoV ORF3b, which is a 154 amino acid ORF whose various Sarbecovirus orthologs are truncated by numerous in-frame stop codons. Its start codon is conserved in all but one of our 44 Sarbecovirus strains, but its stop codon is only present in SARS-CoV-2 and its three closest relatives, and the ORF length is highly variable, so the SARS-CoV-2 form is not (**Extended Data Fig. 6**). Overexpression in a human cell line of the SARS-CoV-2 ORF was found to have anti-IFN-I activity[13]. However, the PhyloCSF score per codon of this truncated ORF is strongly negative (-18.0), it does not overlap any SCEs (**Fig. 2b**), and even among the four closely related strains sharing this stop codon all six substitutions are radical amino acid changes, providing no evidence that this amino acid sequence has been under purifying selection. There is no TRS in the 5’ neighborhood of the ORF3b start codon, and in order for ORF3b to be translated by leaky scanning from the subgenomic RNA for ORF3a, the ribosome would have to bypass eight AUG codons, including several with moderate or strong Kozak context. Finally, ribosome profiling and transcription studies did not find translation of ORF3b or substantial transcription of a subgenomic RNA from which it could be translated[10,15–17].

A different ORF, the 57 amino acid ORF with coordinates 25524-25697, has also been referred to as ORF3b[6,7]. We refer to it as ORF3d to avoid confusion with the 22 amino acid ORF. ORF3d has premature stop codons in all of the other strains of our Sarbecovirus alignment, indicating unambiguously that it does not encode a conserved protein (**Extended Data Fig. 7**), but some authors have suggested it is a *de novo* protein in SARS-CoV-2 based on a sequence analysis method[12] and a statistical test for unexpectedly long overlapping ORFs[11]. On the other hand, analysis of 2,784 SARS-CoV-2 sequences from the GISAID database (Elbe and Buckland-Merrett 2017) found that 17.6% of isolates contain a G to T mutation in position 25563, which introduces a stop codon into the reading frame of ORF3d[7], militating against *de novo* functional translation in SARS-CoV-2. Nor is there convincing experimental evidence that ORF3d is translated. A ribosome profiling study found no evidence of initiation at the start codon of ORF3d in samples collected five hours post infection[10]. Footprints were found at the ORF3d start codon in a reanalysis of footprints from the same study collected 24 hours post infection in samples treated with harringtonine and LTM, which typically would indicate initiating ribosomes[11]; however, the majority of footprints from the 24-hour sample are thought not to be generated by ribosome protection, possibly instead originating from protection by the N protein, and so are not an indication of translation [10]. A subset of ORF3d in the same reading frame, 33 amino acid 3a.iORF2, that begins at a downstream AUG and does not include the nonsense mutation, was one of several overlapping ORF candidates predicted to be translated using ribosome profiling [10]; however, this ORF is not conserved in any other strain and it is unclear how it could be translated: translation by leaky scanning from the ORF3a subgenomic RNA would require the ribosome to bypass 4 earlier AUG codons, and there is no TRS near its start codon that might give rise to a subgenomic RNA specialized for this ORF.

### Supplementary Text S3 Search for other novel protein-coding genes

As discussed in the main text, we computed PhyloCSF scores for all 67 hypothetical non-NCBI-annotated AUG-to-stop locally-maximal SARS-CoV-2 ORFs ≥25 codons long and found that the top scoring candidate, ORF3c, is protein-coding.

Continuing down the sorted list in order of decreasing PhyloCSF score per codon, we found the candidate ORF with the next best score (-2.74) is a 32-codon ORF (26183-26278) that overlaps the 3' end of ORF3a and the 5' end of E (**Supplementary Fig. S4a**) and is orthologous to the 5' end of SARS-CoV ORF3b. Two strains have a frame-shifting 1-base deletion within the ORF, and two others have premature stop codons. None of the substitutions are synonymous. There is high nucleotide-level constraint, but it continues on both sides of the ORF, suggesting it results from something other than translation of the ORF. Overall, this ORF does not show the evolutionary signature of a functional coding sequence. Next in the list is ORF9b which we have discussed elsewhere. Fourth is a 31-codon ORF (3207-3299) overlapping ORF1a, having PhyloCSF score -7.77 (**Supplementary Fig. S4b**). Most of this ORF consists of a 75-nt insertion that is only present in SARS-CoV-2, RaTG13, and CoVZC45, and the start and stop codons are missing in CoVZC45, so this is not a conserved coding sequence. Finally, the fifth-ranked candidate is ORF9c, which we have discussed elsewhere.

The relatively high scores of ORFs 9b and 9c among these 67 hypothetical ORFs are, in part, an artifact of the low density of substitutions throughout N, which they both overlap. This low density, which is found even in the parts of N that are not in ORFs 9b or 9c, decreases the number of substitutions available to PhyloCSF for distinguishing its coding and noncoding evolutionary models, which compresses the PhyloCSF score towards 0, resulting in a better rank among the negative scores. If we compensate for this by dividing by the average number of substitutions per site in the ORF, ORFs 9b and 9c, while still in the top half, move down to the 92nd and 83rd percentile among the 67 ORFs considered, whereas ORF3c remains the best scoring-candidate (**Extended Data Fig. 8**).

To search for additional novel protein-coding regions, we relaxed our criteria to include ORFs with at least 10 codons, allow near-cognate start codons, and allow ORFs contained within another ORF in the same frame. The most promising were two non-canonical ORFs in the 5'-UTR with slightly positive PhyloCSF score, 14-codon CUG-initiated NC_045512.2:92-133 and 31-codon AGG initiated NC_045512.2:158-250. However, neither are among the translated ORF candidates identified by ribosome profiling[10], and the evolutionary evidence is not strong enough to consider it likely that such short non-canonical ORFs generate proteins. Because it has been conjectured that translation might occur on the large number of negative-strand genomic and subgenomic RNAs that are intermediates in viral gene expression and replication in positive-strand RNA viruses [56,57], we also scored ORFs on the negative strand, but again found no convincing candidates. Supplementary Table S4 contains the complete list of ORFs, with scores and other pertinent information.

### Supplementary Text S4 Experimental evidence supporting each ORF

Among previously ambiguous cases, our new reference gene set includes 3c and 9b, and excludes 3b, 3d, 10, and 9c. These inclusion/exclusion decisions are supported by the experimental evidence. Direct RNA sequencing detected little or no subgenomic RNA for the excluded ORFs[15–18], whereas included ORFs 3c and 9b are near the 5' ends of ORFs 3a and N, where they could be translated via leaky scanning from the corresponding subgenomic RNAs. Translation of 3c and 9b, but not 3d, 3b, or 9c, was detected in ribosome profiling experiments[10], and footprints in ORF10 appear to support non-functional translation of overlapping ORFs instead. Peptides supporting translation of 9b but not 10 or 9c were detected in proteomics experiments [15,36] (3b, 3c, and 3d were not included in the search space). Finally, there is a wealth of experimental evidence supporting translation of all of the other named and unnamed ORFs[10,14–18]. This information is summarized in Supplementary Table S2.

### Supplementary Text S5 Effect of reference gene set on variant classification

Having correct gene annotations is critical for determining the effects of variants because the first step in variant classification is understanding how each variant affects protein sequence. Using our reference gene set instead of previous annotations improves the classification of many variants.

In particular, we found that seven variants within ORF3c (T25473C, T25476C, G25494T, G25500A, G25500T, C25539T, and C25572T) that were classified by Nextstrain as synonymous based on their predicted effect on ORF3a, but that we now recognize disrupt amino acids in the ORF3c protein, and three variants (C25493T, G25563T, and A25575C) induce a more radical amino acid change in ORF3c than in ORF3a. Similarly, seven variants within ORF9b (C28291T, T28297C, C28315T, C28369T, G28378T, C28432T, C28519T) were considered synonymous changes in N according to the NCBI annotations, but disrupt amino acids in ORF9b, and two others (G28300T, G28357T) induce a more radical amino acid change in ORF9b than in N. Nextstrain also classified ten variants as amino acid changes in ORF10 that we now recognize are non-coding variants, and five variants as amino acid changes in ORF9c that are synonymous in N and therefore do not change any protein.

Supplementary Table S3 includes Nextstrain variants annotated with respect to the current NCBI reference gene annotations and also with respect to our proposed new reference annotations in. The INFO field also includes Nextstrain's classification according to UniProt annotations.

### Supplementary Text S6 Explanations for within-/cross-strains deviation in nsp3/S1

We propose several possible explanations for the depletion of amino-acid-changing mutations in S1 and nsp3 relative to what would be expected from the overall trend given their high inter-strain evolutionary rates. These differences might indicate:

- Recent changes in their mutation rates, or recent changes in their selective pressures, which could be inherent to SARS-CoV-2 independently of the current pandemic, or could stem from different pressures acting on these genes during pandemic expansion in human hosts, relative to stable spread of Sarbecoviruses in bat populations
- These differences may also reflect general properties in an adaptation-expansion cycle of coronaviruses, with S1 and nsp3 initially undergoing rapid evolution during adaptation to a new host, followed by a period in which purifying selection suppresses further variation. In this scenario, the smaller-than-expected number of observed mutations in the current pandemic would stem from pre-adaptation of S1 and nsp3, either through transmission in non-human animal hosts with a similar ACE2 receptor, or through undetected transmission in humans prior to when the initial sample for the reference genome was obtained in December

2019 (Wu et al.), thus requiring relatively fewer human-adaptive mutations compared to other genes whose biological functions would adapt to human hosts only later (noting however, that only a subset of mutations in the current pandemic are likely adaptive).

- The frequent recombination observed in S1 suggests an alternative explanation. It is possible that recombination events into our selected 44-strain phylogeny from more distant relatives could have increased the number of inter-strain differences in these genes. In that case, the paucity of mutations in S1 and nsp3 relative to their inter-strain differences might be due to inflation of the latter rather than deflation of the former. However, we note that the amount of distant recombination needed to account for the large discrepancy observed might be implausibly large.
- Lastly, inter-strain differences reflect selective pressures that have acted over evolutionary time scales in which even mildly deleterious mutations are excluded, while within-strain differences reflect smaller evolutionary time scales over which only strongly deleterious mutations are excluded. Thus, the discrepancy observed could also result if the fraction of all possible deleterious amino acid changes of S1 and nsp3 that are strongly deleterious rather than mildly deleterious is sufficiently larger than that fraction for other proteins.

### Supplementary Text S7 Enriched and depleted clusters of missense SNVs

Other than the Nucleocapsid region that we have already discussed, there are no regions in the genome in which variants disrupting conserved amino acid residues are significantly denser than would be expected by chance, given the total number of such variants in each gene. Nor are there any regions that are significantly depleted for such variants, which would have indicated regions in which constraint in the Sarbecovirus clade has continued particularly strongly in the SARS-CoV-2 population. The regions that are most depleted (though not statistically significantly) have coordinates 7400-7840 in nsp3 with no missense variants among 103 conserved amino acids and 24437-24748 in S2 with no missense variants among 99 conserved amino acids (p=0.072 and p=0.093, respectively, without any correction for multiple region lengths searched) (**Supplementary Fig. S5**).
