## Supplementary material for "SARS-CoV-2 gene content and COVID-19 mutation impact by comparing 44 Sarbecovirus genomes": Summary Of Supplementary Data

### Supplementary Information:

- **Supplementary Data.docx:** Supplementary Figures S1-S5 and Supplementary Text S1-S8.
- **Supplementary Table S1.txt:** Tab-delimited table with one row for each of 44 Sarbecovirus strains used.
- **Supplementary Table S2.txt**: Tab-delimited table with information on all confirmed or proposed ORFs and mature proteins of SARS-CoV-2.
- **Supplementary Table S3.txt:** Tab-delimited table with information about each of the single nucleotide variants used in this study
- **Supplementary Table S4:.txt** Tab-delimited table listing open reading frames searched for novel coding regions.
- **PhyloCSFGenes.bed.txt:** Our proposed new reference gene set for SARS-CoV-2 in BED format.
- **PhyloCSFRejectedGenes.bed.txt:** The genes others have proposed that we rejected in BED format.
- **44Sarbecoviruses.nh.txt**: Whole-genome phylogenetic tree in Newick format.
- **44Sarbecoviruses.fa.txt**: Whole-genome alignment of 44-Sarbecovirus genomes in Fasta format.
- **nextstrain_ncov_global_metadata.txt**: List of authors who contributed genomes to GISAID that were used by Nextstrain and UCSC to produce the list of SNVs.
