## Supplementary material for "SARS-CoV-2 gene content and COVID-19 mutation impact by comparing 44 Sarbecovirus genomes": CodAlignView PDFs For All Genes: CodAlignView nsp11.pdf

| NC_045512_SARS_CoV_2_Wuhan_Hu_1_aa | E | P | M | L | Q | S | A | D | A | Q | S | F | L | N | G | F | A | V | * | V | Q | P | V |
|---|---|---|---|---|---|---|---|---|---|---|---|---|---|---|---|---|---|---|---|---|---|---|---|
| ORFs and Mature Proteins | | <- | nsp 10 | | ->/ | \|<- | ns p11 | | -> | | | | | | <- | nsp 11 | | ->\| | | | | | |
| NC_045512_SARS_CoV_2_Wuhan_Hu_1 | GAA | CCC | ATG | CTT | CAG | TCA | GCT | GAT | GCA | CAA | TCG | TTT | TTA | AAC | GGG | TTT | GCG | GTG | TAA | GTG | CAG | CCC | GTC |
| MN996532_Bat_CoV_RaTG13 | GAA | CCC | ATG | CTT | CAG | TCA | GCT | GAT | GCA | CAA | TCG | TTT | TTA | AAC | GGG | TTT | GCG | GTG | TAA | GTG | CAG | CCC | GTC |
| MG772933_Bat_SARS_like_CoV_bat_SL_CoVZC45 | GAA | CCC | ATG | ATG | CAG | TCT | GCG | GAT | GCG | TCA | ACG | TTT | TTA | AAC | GGG | TTT | GCG | GTG | TAA | GTG | CGG | CCC | GTC |
| MG772934_Bat_SARS_like_CoV_bat_SL_CoVZXC21 | GAA | CCC | ATG | ATG | CAG | TCT | GCG | GAC | GCG | TCA | ACG | TTT | TTA | AAC | GGG | TTT | GCG | GTG | TAA | GTG | CAG | CCC | GTC |
| NC_004718_SARS_CoV | GAA | CCC | TTG | ATG | CAG | TCT | GCG | GAT | GCA | TCA | ACG | TTT | TTA | AAC | GGG | TTT | GCG | GTG | TAA | GTG | CAG | CCC | GTC |
| KT444582_SARS_like_CoV_WIV16 | GAA | CCC | ATG | ATG | CAG | TCT | GCG | GAT | GCG | TCA | ACG | TTT | TTA | AAC | GGG | TTT | GCG | GTG | TAA | GTG | CAG | CCC | GTC |
| KY417146_Bat_SARS_like_CoV_Rs4231 | GAA | CCC | ATG | ATG | CAG | TCT | GCG | GAT | GCG | TCA | ACG | TTT | TTA | AAC | GGG | TTT | GCG | GTG | TAA | GTG | CAG | CCC | GTC |
| MK211376_CoV_BtRs_BetaCoV_YN2018B | GAA | CCC | ATG | ATG | CAG | TCT | GCG | GAT | GCG | TCA | ACG | TTT | TTA | AAC | GGG | TTT | GCG | GTG | TAA | GTG | CAG | CCC | GTC |
| KY417151_Bat_SARS_like_CoV_Rs7327 | GAA | CCC | ATG | ATG | CAG | TCT | GCG | GAT | GCG | TCA | ACG | TTT | TTA | AAC | GGG | TTT | GCG | GTG | TAA | GTG | CAG | CCC | GTC |
| KY417152_Bat_SARS_like_CoV_Rs9401 | GAA | CCC | ATG | ATG | CAG | TCT | GCG | GAT | GCG | TCA | ACG | TTT | TTA | AAC | GGG | TTT | GCG | GTG | TAA | GTG | CAG | CCC | GTC |
| KY417144_Bat_SARS_like_CoV_Rs4084 | GAA | CCC | ATG | ATG | CAG | TCT | GCG | GAT | GCG | TCA | ACG | TTT | TTA | AAC | GGG | TTT | GCG | GTG | TAA | GTG | CAG | CCC | GTC |
| KF367457_Bat_SARS_like_CoV_WIV1 | GAA | CCC | ATG | ATG | CAG | TCT | GCG | GAT | GCG | TCA | ACG | TTT | TTA | AAC | GGG | TTT | GCG | GTG | TAA | GTG | CAG | CCC | GTC |
| KU973692_UNVERIFIED_SARS_related_CoV_F46 | GAA | CCC | ATG | ATG | CAA | TCT | GCG | GAT | GCG | TCA | ACG | TTT | TTA | AAC | GGG | TTT | GCG | GTG | TAA | GTG | CAG | CCC | GTC |
| KY417145_Bat_SARS_like_CoV_Rf4092 | GAA | CCC | ATG | ATG | CAG | TCA | GCG | GAT | GCG | TCA | ACG | TTT | TTA | AAC | GGG | TTT | GCG | GTG | TAA | GTG | CAG | CCC | GTC |
| KJ473816_BtRs_BetaCoV_YN2013 | GAA | CCC | ATG | ATG | CAG | TCT | GCG | GAT | GCG | TCA | GCG | TTT | TTA | AAC | GGG | TTT | GCG | GTG | TAA | GTG | CAG | CCC | GTC |
| KY770858_Bat_CoV_Anlong_103 | GAA | CCC | ATG | ATG | CAG | TCT | GCG | GAT | GCG | TCA | ACG | TTT | TTA | AAC | GGG | TTT | GCG | GTG | TAA | GTG | CAG | CCC | GTC |
| KY417143_Bat_SARS_like_CoV_Rs4081 | GAA | CCC | ATG | ATG | CAG | TCT | GCG | GAT | GCG | TCA | ACG | TTT | TTA | AAC | GGG | TTT | GCG | GTG | TAA | GTG | CAG | CCC | GTC |
| KY417149_Bat_SARS_like_CoV_Rs4255 | GAA | CCC | ATG | ATG | CAG | TCT | GCA | GAT | GCG | TCA | ACG | TTT | TTA | AAC | GGG | TTT | GCG | GTG | TAA | GTG | CAG | CCC | GTC |
| MK211378_CoV_BtRs_BetaCoV_YN2018D | GAA | CCC | ATG | ATG | CAG | TCT | GCG | GAT | GCG | TCA | ACG | TTT | TTA | AAC | GGG | TTT | GCG | GTG | TAA | GTG | CAG | CCC | GTC |
| FJ588686_Bat_SARS_CoV_Rs672_2006 | GAA | CCC | ATG | ATG | CAG | TCT | GCG | GAC | GCG | TCA | ACG | TTT | TTA | AAC | GGG | TTT | GCG | GTG | TAA | GTG | CAG | CCC | GTC |
| MK211377_CoV_BtRs_BetaCoV_YN2018C | GAA | CCC | ATG | ATG | CAG | TCT | GCG | GAT | GCG | TCA | ACG | TTT | TTA | AAC | GGG | TTT | GCG | GTG | TAA | GTG | CAG | CCC | GTC |
| KY417142_Bat_SARS_like_CoV_As6526 | GAA | CCC | ATG | ATG | CAG | TCT | GCG | GAT | GCG | TCA | ACG | TTT | TTA | AAC | GGG | TTT | GCG | GTG | TAA | GTG | CAG | CCC | GTC |
| KY417147_Bat_SARS_like_CoV_Rs4237 | GAA | CCC | ATG | ATG | CAG | TCT | GCG | GAT | GCG | TCA | ACG | TTT | TTA | AAC | GGG | TTT | GCG | GTG | TAA | GTG | CAG | CCC | GTC |
| KY417148_Bat_SARS_like_CoV_Rs4247 | GAA | CCC | ATG | ATG | CAG | TCT | GCG | GAT | GCG | TCA | ACG | TTT | TTA | AAC | GGG | TTT | GCG | GTG | TAA | GTG | CAG | CCC | GTC |
| MK211375_CoV_BtRs_BetaCoV_YN2018A | GAA | CCC | ATG | ATG | CAG | TCT | GCG | GAT | GCG | TCA | ACG | TTT | TTA | AAC | GGG | TTT | GCG | GTG | TAA | GTG | CAG | CCC | GTC |
| DQ071615_Bat_SARS_CoV_Rp3 | GAA | CCC | ATG | ATG | CAG | TCT | GCG | GAT | GCG | TCA | ACG | TTT | TTA | AAC | GGG | TTT | GCG | GTG | TAA | GTG | CAG | CCC | GTC |
| KP886808_Bat_SARS_like_CoV_YNLF_31C | GAA | CCC | ATG | ATG | CAG | TCT | GCG | GAT | GCG | TCA | ACG | TTT | TTA | AAC | GGG | TTT | GCG | GTG | TAA | GTG | CAG | CCC | GTC |
| KJ473815_BtRs_BetaCoV_GX2013 | GAA | CCC | ATG | ATG | CAG | TCT | GCG | GAT | GCG | TCA | ACG | TTT | TTA | AAC | GGG | TTT | GCG | GTG | TAA | GTG | CAG | CCC | GTC |
| KF569996_Rhinolophus_affinis_CoV_LYRa11 | GAA | TCC | ATG | ATG | CAG | TCT | GAG | GAC | GCG | TCA | ACT | TTT | TTA | AAC | GGG | TTT | GCG | GTG | TAA | GTG | CAG | CCC | GTC |
| JX993988_Bat_CoV_Cp_Yunnan2011 | GAA | CCC | ATG | ATG | CAG | TCT | GCG | GAC | GCG | TCA | ACG | TTT | TTA | AAC | GGG | TTT | GCG | GTG | TAA | GTG | CAG | CCC | GTC |
| MK211374_CoV_BtRl_BetaCoV_SC2018 | GAA | CCC | ATG | ATG | CAG | TCT | GCG | GAT | GCG | TCA | ACG | TTT | TTA | AAC | GGG | TTT | GCG | GTG | TAA | GTG | CAG | CCC | GTC |
| KJ473814_BtRs_BetaCoV_HuB2013 | GAA | CCC | ATG | ATG | CAG | TCT | GCT | GAC | GCA | TCA | ACG | TTT | TTA | AAC | GGG | TTT | GCG | GTG | TAA | GTG | CAG | CCC | GTC |
| DQ412043_Bat_SARS_CoV_Rm1 | GAA | CCC | ATG | ATG | CAG | TCT | GCT | GAC | GCG | TCA | ACG | TTT | TTA | AAC | GGG | TTT | GCG | GTG | TAA | GTG | CGG | CCC | GTC |
| KY938558_Bat_CoV_strain_16BO133 | GAA | CCC | ATG | ATG | CAG | TCT | GCG | GAT | GCG | TCA | CCG | TTT | TTA | AAC | GGG | TTT | GCG | GTG | TAA | GTG | CAG | CCC | GTC |
| DQ412042_Bat_SARS_CoV_Rf1 | GAA | CCC | ATG | ATG | CAG | TCG | GCG | GAT | GCG | TCA | CCG | TTT | TTA | AAC | GGG | TTT | GCG | GTG | TAA | GTG | CAG | CCC | GTC |
| DQ648856_Bat_CoV_BtCoV_273_2005 | GAA | CCC | ATG | ATG | CAG | TCG | GCG | GAT | GCG | TCA | CCG | TTT | TTA | AAC | GGG | TTT | GCG | GTG | TAA | GTG | CAG | CCC | GTC |
| KJ473812_BtRf_BetaCoV_HeB2013 | GAA | CCC | ATG | ATG | CAG | TCT | GCG | GAT | GCG | TCA | GCG | TTT | TTA | AAC | GGG | TTT | GCG | GTG | TAA | GTG | CAG | CCC | GTC |
| KY770860_Bat_CoV_Jiyuan_84 | GAA | CCC | ATG | ATG | CAG | TCT | GCG | GAT | GCG | TCA | GCG | TTT | TTA | AAC | GGG | TTT | GCG | GTG | TAA | GTG | CAG | CCC | GTC |
| JX993987_Bat_CoV_Rp_Shaanxi2011 | GAA | CCC | ATG | ATG | CAG | TCT | GCT | GAC | GCA | TCA | ACG | TTT | TTA | AAC | GGG | TTT | GCG | GTG | TAA | GTG | CAG | CCC | GTC |
| GQ153542_Bat_SARS_CoV_HKU3_7 | GAA | CCC | ATG | ATG | CAG | TCT | GCG | GAC | GCG | TCA | ACG | TTT | TTA | AAC | GGG | TTT | GCG | GTG | TAA | GTG | CGG | CCC | GTC |
| DQ022305_Bat_SARS_CoV_HKU3_1 | GAA | CCC | ATG | ATG | CAG | TCT | GCG | GAT | GCG | TCA | ACG | TTT | TTA | AAC | GGG | TTT | GCG | GTG | TAA | GTG | CGG | CCC | GTC |
| GQ153547_Bat_SARS_CoV_HKU3_12 | GAA | CCC | ATG | ATG | CAG | TCT | GCG | GAT | GCG | TCA | ACG | TTT | TTA | AAC | GGG | TTT | GCG | GTG | TAA | GTG | CGG | CCC | GTC |
| KY352407_SARS_related_CoV_strain_BtKY72 | GAA | CCC | ATG | ATG | CAA | TCA | TCA | GAT | GCG | TCA | ACG | TTT | TTA | AAC | GGG | TTT | GCG | GTG | TAA | GTG | CAG | CCC | GTC |
| NC_014470_Bat_CoV_BM48_31_BGR_2008 | GAG | CCC | GTG | ATG | CAG | GCA | GCT | GAT | GCC | CCA | GCG | TTT | TTA | AAC | GGG | TTT | GCG | GTG | TAA | GTG | CGG | CCC | GTC |

nsp11
