## Supplementary figures and images for "SARS-CoV-2 gene content and COVID-19 mutation impact by comparing 44 Sarbecovirus genomes"

### CodAlignView 1a.pdf

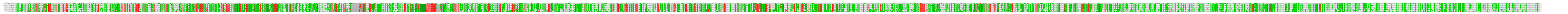

### CodAlignView 1ab.pdf

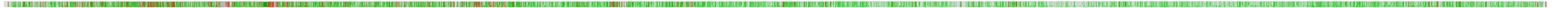

### CodAlignView 3a.pdf

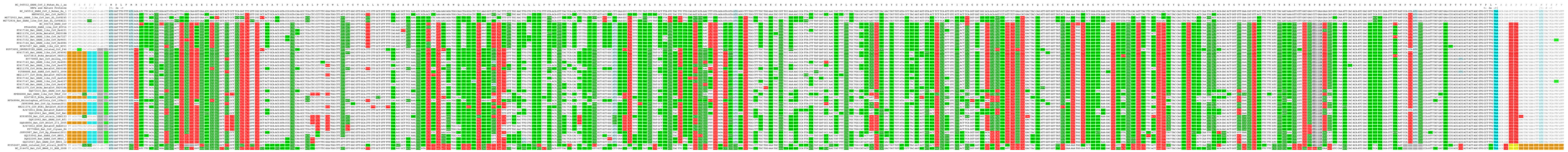

3a

### CodAlignView 6.pdf

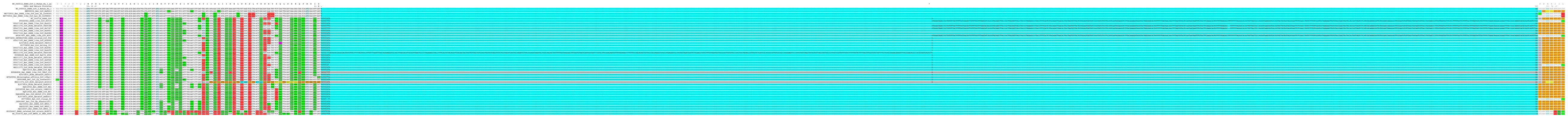

### CodAlignView 7a.pdf

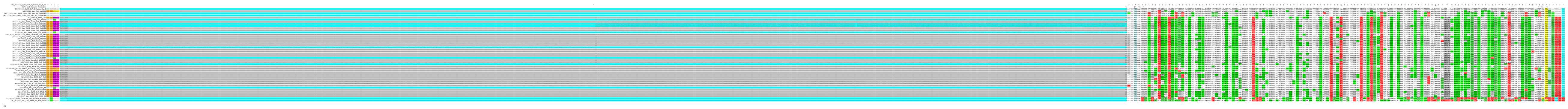

### CodAlignView 7b.pdf

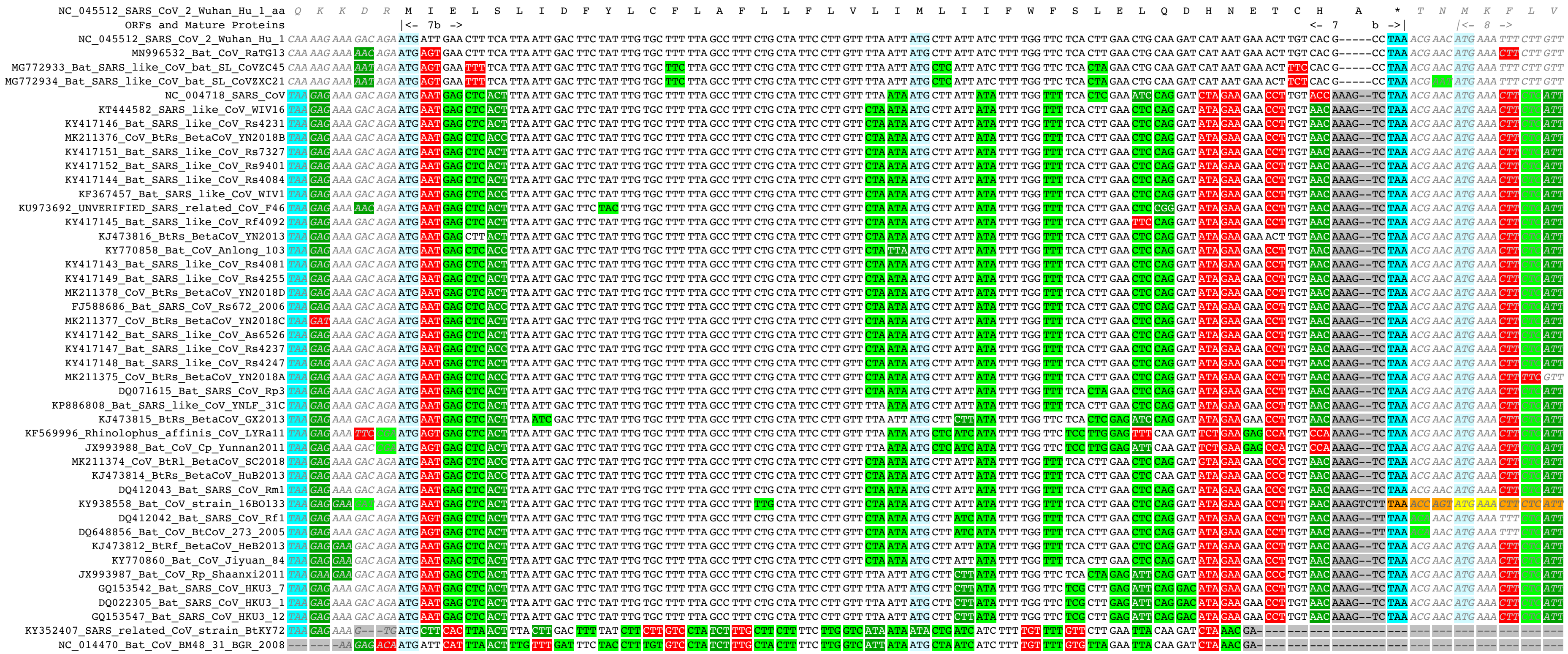

7b

### CodAlignView 8.pdf

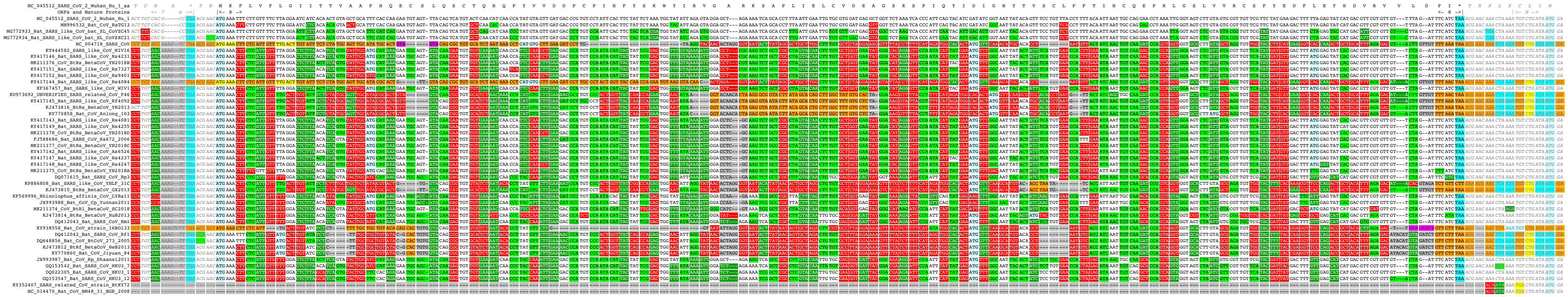

### CodAlignView 9b.pdf

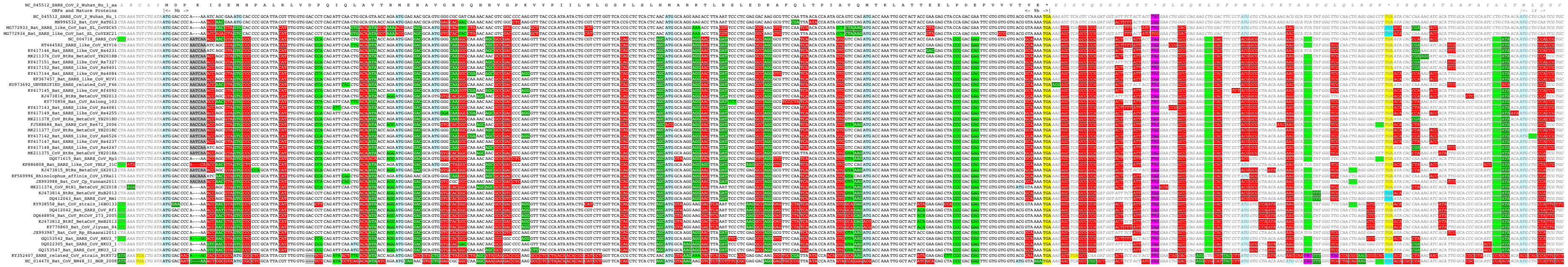

9b

### CodAlignView 9c.pdf

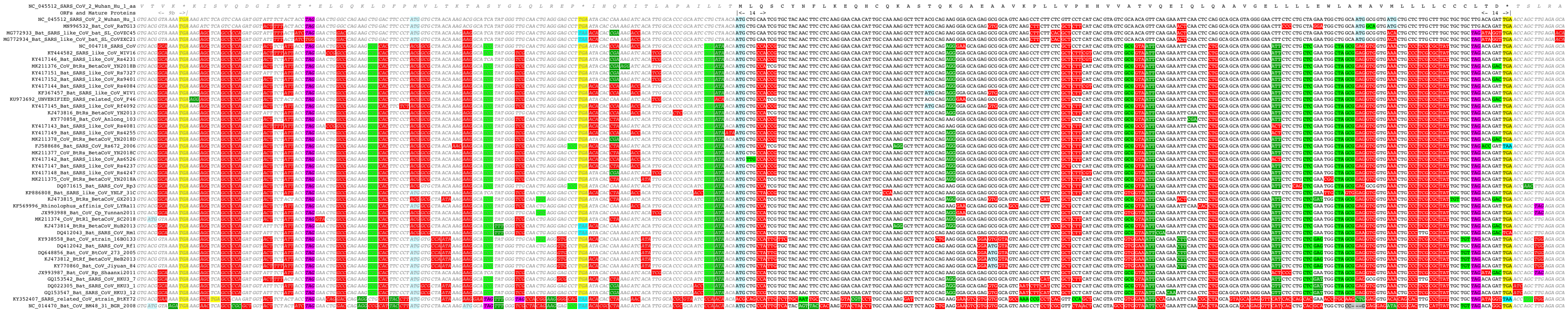

### CodAlignView 10.pdf

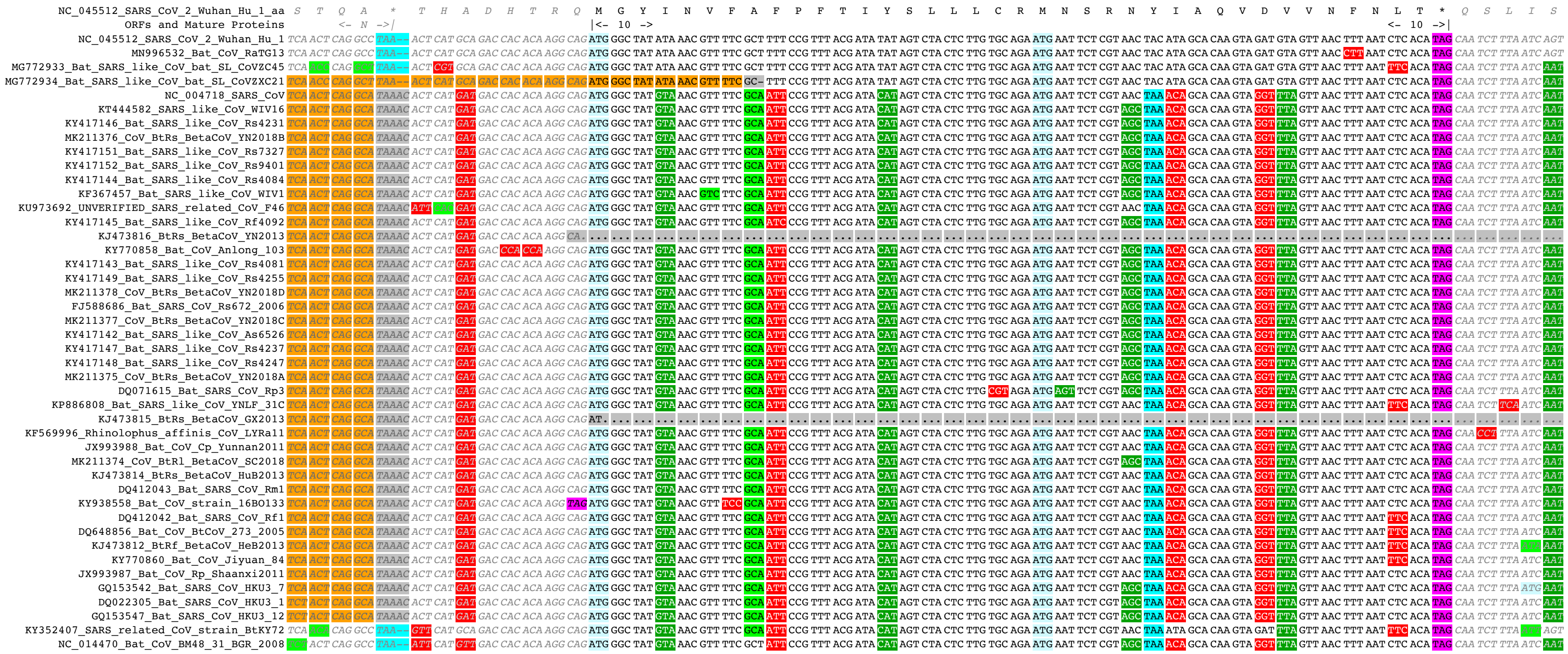

### CodAlignView E.pdf

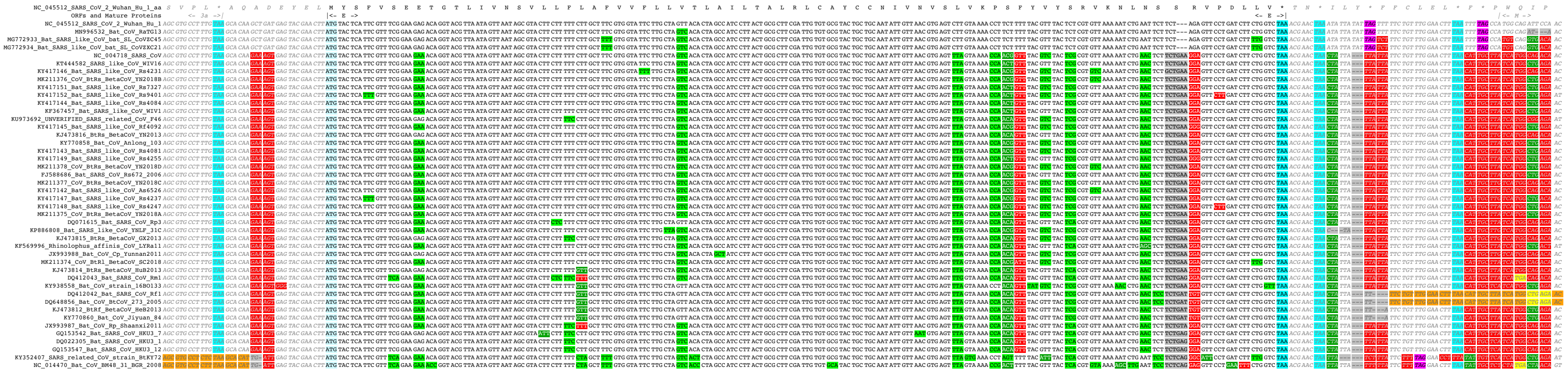


E

### CodAlignView M.pdf

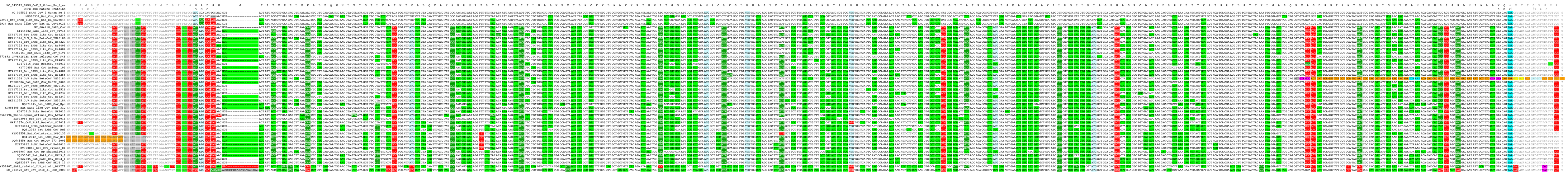

M

### CodAlignView N.pdf

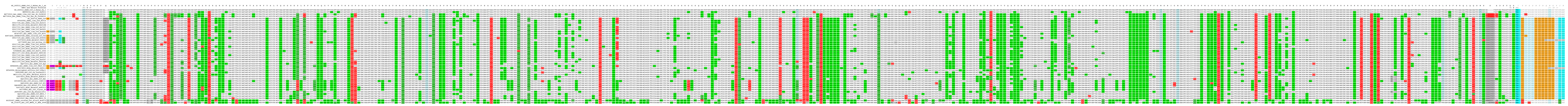

### CodAlignView nsp1.pdf

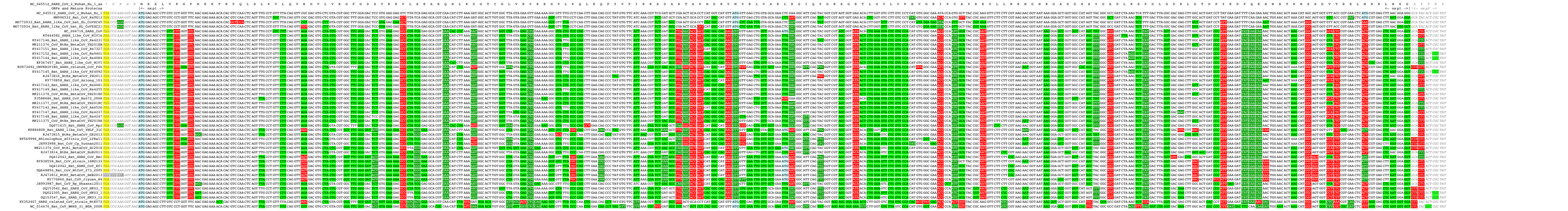

nsp1

### CodAlignView nsp2.pdf

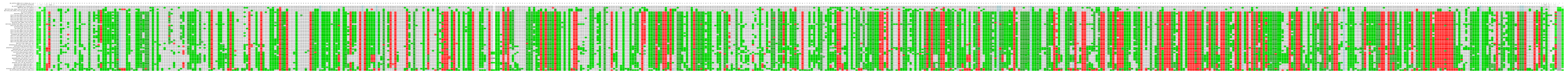

nsp2

### CodAlignView nsp3.pdf

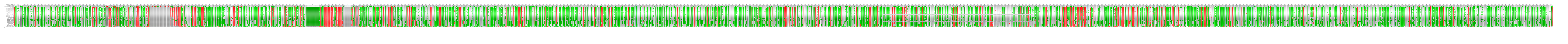

### CodAlignView nsp4.pdf

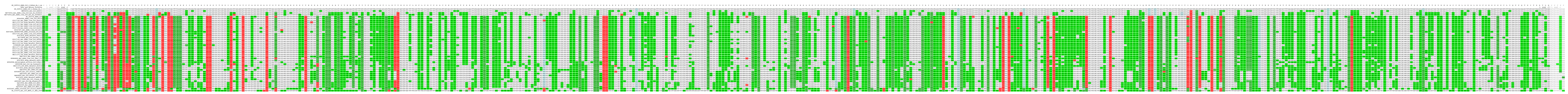

nsp4

### CodAlignView nsp5.pdf

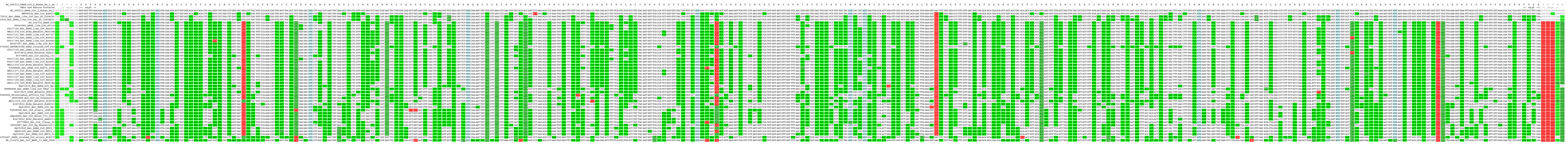

nsp5

### CodAlignView nsp6.pdf

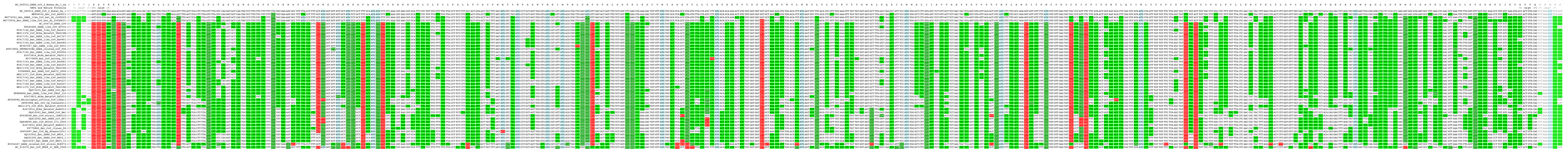

nsp6

### CodAlignView nsp7.pdf

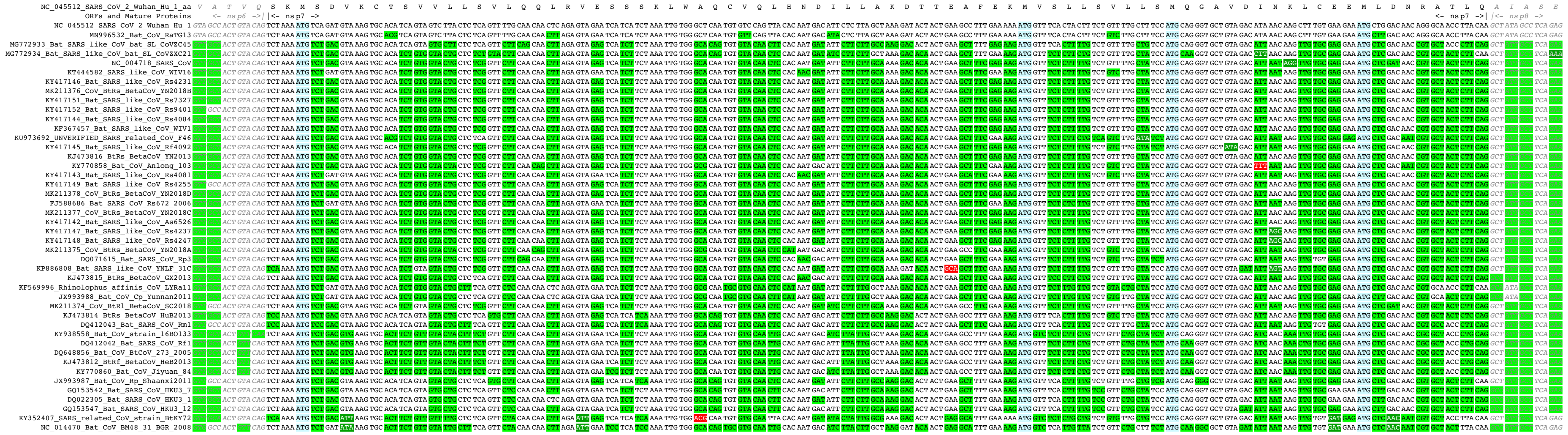

nsp7

### CodAlignView nsp8.pdf

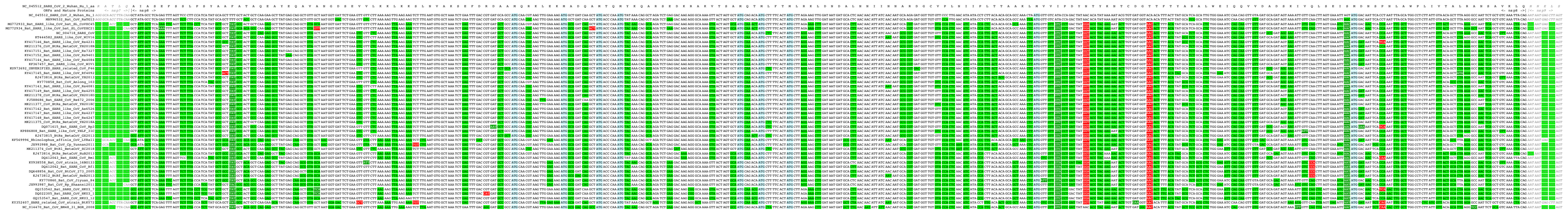

### CodAlignView nsp9.pdf

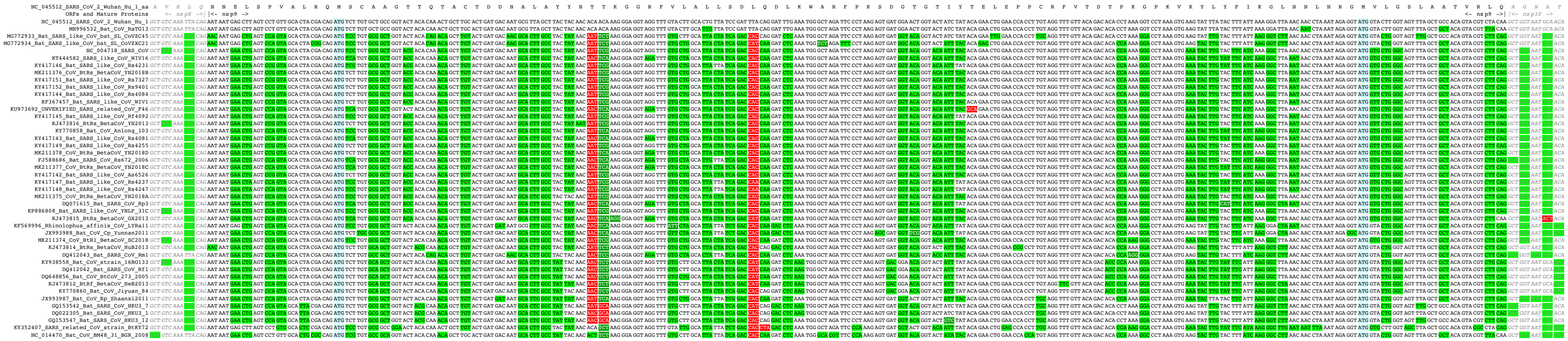

nsp9

### CodAlignView nsp10.pdf

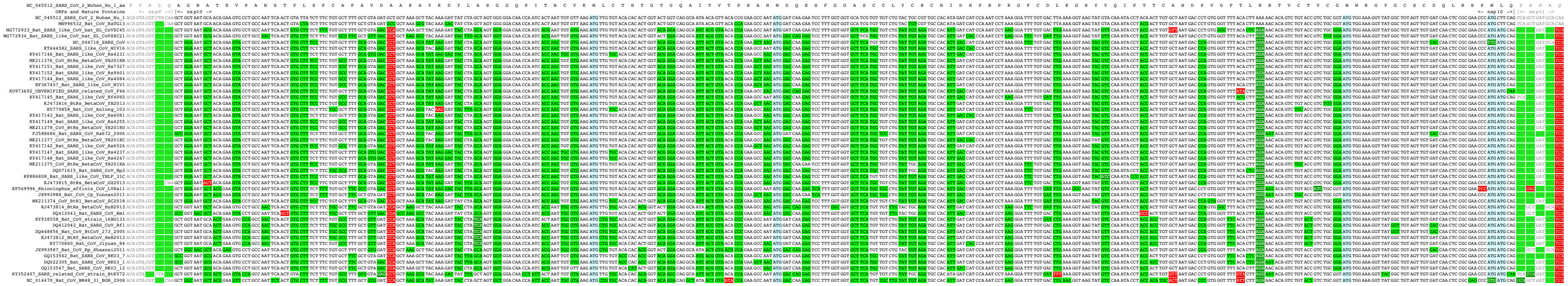

nsp10

### CodAlignView nsp12.pdf

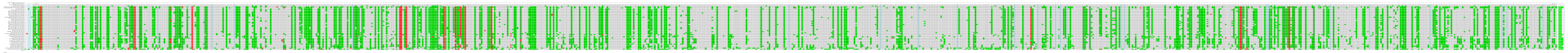

nsp12

### CodAlignView nsp13.pdf

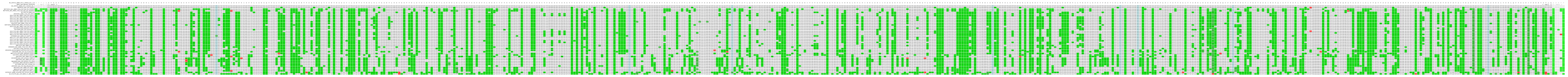

nsp13

### CodAlignView nsp14.pdf

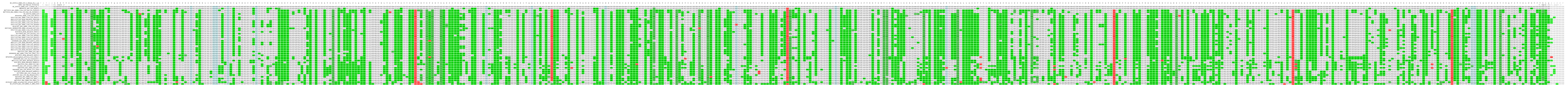

nsp14

### CodAlignView nsp15.pdf

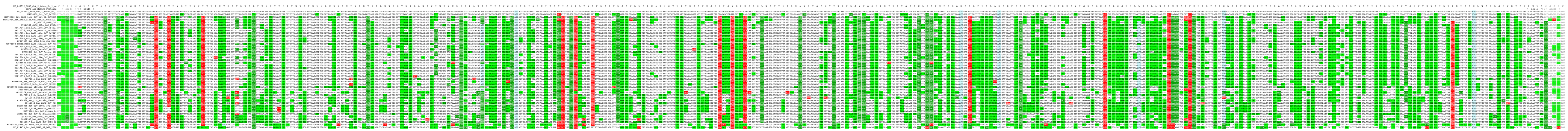

nsp15

### CodAlignView nsp16.pdf

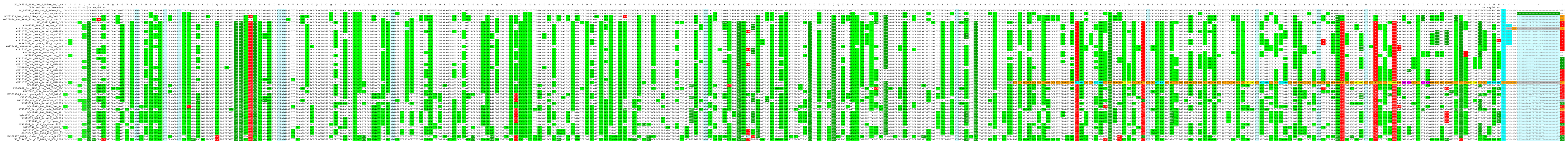

nsp16

### CodAlignView S1.pdf

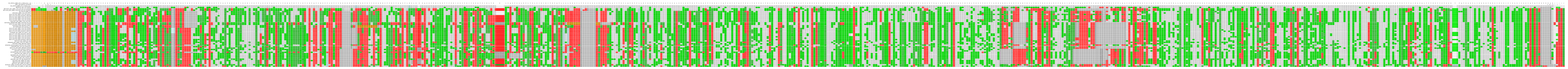

S1

### CodAlignView S2'.pdf

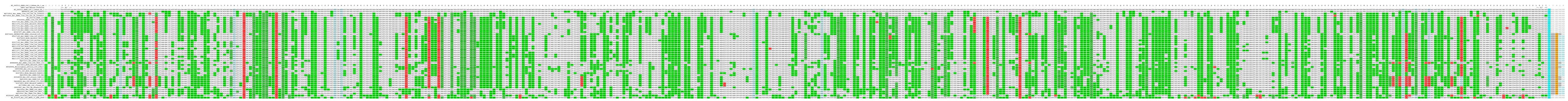

S2'

### CodAlignView S2.pdf

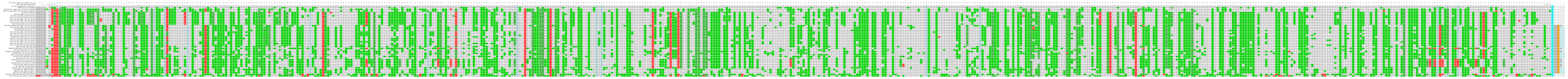

S2

### CodAlignView S.pdf

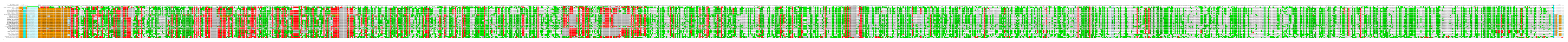
